# Interhemispheric gamma synchrony between parvalbumin interneurons supports behavioral adaptation

**DOI:** 10.1101/784330

**Authors:** Kathleen K.A. Cho, Thomas J. Davidson, Jesse D. Marshall, Mark J. Schnitzer, Vikaas S. Sohal

## Abstract

Organisms must learn novel strategies to adapt to changing environments. Synchrony, which enhances neuronal communication, might create dynamic brain states, facilitating such adaptation. Although synchronization is common in neural systems, its functional significance remains controversial. We studied the role of gamma-frequency (~40 Hz) synchronization, promoted by parvalbumin interneurons, in mice learning multiple new cue-reward associations. Voltage imaging revealed cell type-specific increases of interhemispheric gamma synchrony within prefrontal parvalbumin interneurons, when mice received feedback that previously-learned associations were no longer valid. Disrupting this synchronization by delivering out-of-phase optogenetic stimulation caused mice to perseverate on outdated associations, an effect not reproduced by stimulating in-phase or out-of-phase at other frequencies. Gamma synchrony was specifically required when new associations utilized familiar cues that were previously irrelevant to behavioral outcomes, not when associations involved novel cues, or for reversing previously learned associations. Thus, gamma synchrony is indispensable for reappraising the behavioral salience of external cues.

## Introduction

Adapting to changes in the environment requires the ability to detect when previously learned behavioral strategies are no longer effective, suppress actions based on those outdated strategies, and develop appropriate new ones. The inability to perform these functions is a hallmark of prefrontal cortex dysfunction in schizophrenia, classically measured by paradigms such as the Wisconsin Card Sorting Task (WCST)^1,2^. Critically, many natural behaviors, as well as tasks like the WCST, require organisms to rapidly learn novel strategies that rely on external cues which were previously unimportant or ignored. Dynamic processes within the prefrontal cortex that facilitate this kind of behavioral adaptation remain unknown. Synchrony, particularly in the gamma-frequency range (~30-80 Hz), has been proposed to regulate how neurons interact with each other and their downstream targets^3–13^. Thus, by selectively enhancing interactions between neurons in specific brain regions at particular moments, synchrony could generate dynamic brain states to support behavioral adaptation. In particular, we and others have observed synchronized gamma-frequency activity in the medial prefrontal cortex (mPFC) of rodents during changes in behavior^14–16^. While interneurons, particularly parvalbumin (PV) interneurons, are known to generate synchronized gamma-frequency activity, it remains deeply controversial whether gamma synchronization across brain regions contributes to behavior or simply reflects the increased recruitment of PV interneurons^17,18^.

To address these issues, we explored the role of gamma-frequency synchronization between prefrontal PV interneurons as mice perform a task involving the type of behavioral adaptation outlined above. Variants of this task have previously been characterized by our laboratory and others^14,15,19–22^. On each trial, mice choose between two bowls to find a hidden food reward (Fig. 1a). Each bowl is filled with different odor and texture cues. Mice first form an *initial association* between one type of cue and reward. For example, in an odor-based rule, one odor is consistently located in the rewarded bowl, the other odor is consistently in the unrewarded bowl, and the location of each texture (in the rewarded vs. unrewarded bowl) varies randomly from trial to trial. Then mice must learn a “rule shift” from odor to texture or vice versa. During a rule shift, a familiar *but previously irrelevant* cue becomes associated with reward. This is distinguished from a “rule reversal” in which the type of rule – odor or texture – does not change, but the cue which previously predicted the absence of reward now becomes rewarded (Fig. 1b). Unlike tasks requiring well-trained mice to switch between previously learned behaviors (e.g., pressing a right or left lever), in this task, mice form new associations using familiar cues that were either previously irrelevant (rule shift) or predictive (rule reversal) with respect to behavioral outcomes.

**Fig. 1:**
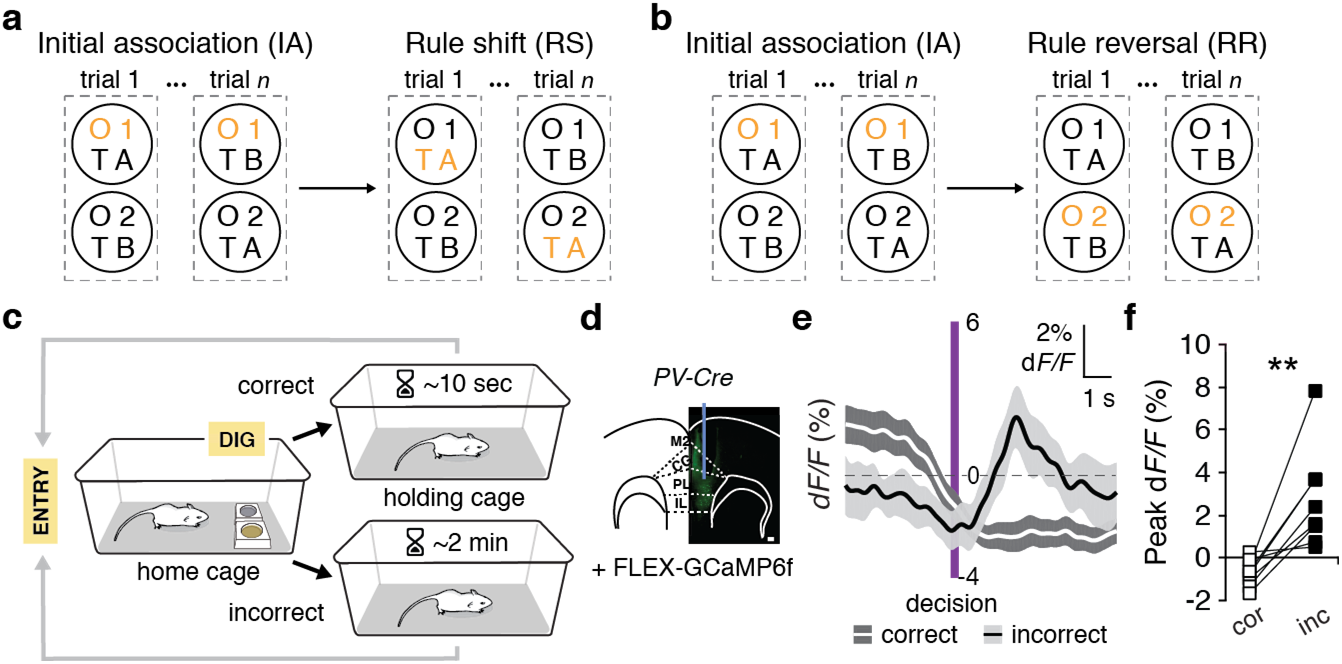
Prefrontal PV interneurons are recruited after errors during rule shifts. **a**, Rule-shift task schematic. On each trial, a mouse chooses one of two bowls, each scented with a different odor (O1 or O2) and filled with a different textured digging medium (TA or TB), to find a food reward. Mice first learn an initial association (IA) between one of these cues (e.g., odor O1) and food reward (the cue associated with reward is indicated in orange). Once mice reach the learning criterion (8/10 consecutive trials correct), this association undergoes an extra-dimensional rule shift (RS; e.g., from O1 to TA). **b**, Rule-reversal task schematic. Mice learn an initial association (IA) between one cue (e.g., odor O1) and food reward (the rewarded cue is indicated in orange). Once mice reach the learning criterion, this association undergoes an intra-dimensional rule reversal (RR), e.g., from O1 to O2. **c**, Trial timeline. A mouse begins each trial by entering the home cage, then makes a decision, indicated by digging in one bowl. If the mouse is correct, food reward is consumed. The mouse is then transferred to the holding cage until the next trial. The intertrial interval is longer after incorrect choices. **d**, Representative image showing mPFC FLEX-GCaMP6f expression in a *PV-Cre* mouse (scale bar, 100 μm). **e**, Averaged PV interneuron photometry signal (*dF/F*), aligned to the time of dig, which indicates a decision, for correct (white line) vs. incorrect trials (black line; *n* = 8). **f**, Peak *dF/F* during the 4 sec following the decision. Signals are significantly higher on incorrect than correct trials (two-tailed, paired *t*-test; *t_(7)_* = 3.93, \*\**P* = 0.006). Data are shown as means (e); shading (e) denote s.e.m.

Several findings from our lab and others suggest that learning during rule shifts, but not during rule reversals, depends on interneuron-generated rhythmic activity in the mPFC. First, mPFC lesions disrupt learning during rule shifts but not during rule reversals^19^. Second, inhibiting GABAergic interneurons in mPFC similarly disrupts learning during rule shifts but not during initial associations or rule reversals^14^. Third, mutant mice with abnormal PV interneurons and deficient task-evoked gamma power in mPFC also have impaired learning during rule shifts but not during initial associations or rule reversals^14^. In fact, in these mutant mice, we previously showed that stimulating prefrontal interneurons at gamma-frequencies (40 or 60 Hz) can normalize learning during rule shifts^14^. This shows that inducing synchronized gamma-frequency activity in prefrontal cortex can affect behavior under pathological conditions. However, this study did not address whether synchronization across brain regions contributes to normal behavior. In fact, similar to our previous work, several other studies have shown that optogenetically stimulating prefrontal interneurons at gamma frequencies can improve sensory detection, attention, and social behavior^22–24^. However, none of these have directly addressed two fundamental, outstanding questions: first, do these behavioral effects require the synchronization of activity *across brain regions*, or is enhancing rhythmic inhibition within a local circuit sufficient? Second, even if artificially increasing synchrony can improve behavior, is *naturally-occurring synchrony necessary for normal behavior?*

Here we explored how interhemispheric gamma synchrony between prefrontal PV interneurons contributes to learning new associations during a rule shift. Developing new analyses based on genetically encoded voltage indicators, we find that gamma-frequency synchrony between PV interneurons in the left and right mPFC increases when mice learn new associations during rule shifts. However, gamma synchrony does not increase during learning of initial associations or rule reversals. Using optogenetics, we then find that perturbing this naturally-occurring synchronization, by delivering weak gamma-frequency optogenetic stimulation to PV interneurons *out-of-phase* between the left and right mPFC, disrupts learning during a rule shift. The same stimulation has no effect when delivered *inphase*. This dissociation between the effects of in vs. out-of-phase stimulation disambiguates the role of interhemispheric synchrony from that of rhythmic inhibition within a local circuit. Furthermore, optogenetically perturbing interhemispheric gamma synchrony does not affect learning during initial associations or rule reversals. Thus, there is a 1:1 correspondence between whether a type of learning normally elicits increased gamma synchrony, and whether it is disrupted by perturbing that synchrony.

Finally, to independently validate this relationship between gamma synchrony and learning during rule shifts, we studied mutant mice that have impaired learning during rule shifts (but not during initial associations or rule reversals). We find impaired gamma synchrony in these mice specifically during rule shifts; restoring their gamma synchrony either optogenetically or pharmacologically rescues learning. These results show that gamma synchrony across the hemispheres is normally recruited by and necessary for a specific form of learning, which involves developing new strategies using familiar cues that were previously unimportant.

## Results

### PV interneurons are recruited after errors during rule shifts

Previous work has shown that optogenetically inhibiting prefrontal PV interneurons disrupts learning during rule shifts^21^. Therefore, as a first step, we used bulk calcium imaging^25–26^ (fiber photometry) to explore when PV interneurons are normally recruited during rule shifts (Fig. 1d–f; Supplementary Fig. 1). We targeted PV interneurons by injecting AAV1-Syn-FLEX-GCaMP6f^27^ (Methods) unilaterally into the mPFC (near the prelimbic/infralimbic border) of *PV-Cre* mice and implanted a multimode optical fiber to detect population-level changes in fluorescent calcium signals (Fig. 1d). We examined activity time-locked to readily-definable task events: the start of each trial, the time of each decision (indicated by the mouse beginning to dig in one bowl), the end of each trial (when the mouse stopped digging in that bowl), and the intertrial interval (Fig. 1c). On error trials, there was a marked rise in PV interneuron activity following decisions, i.e., when animals failed to receive a reward that would have been expected based on the previously learned association (Fig. 1e,f). In contrast, PV activity did not increase following correct decisions (Fig. 1e,f). Because the rule shift is uncued, the failure to receive an expected reward can serve as an indication that a previously learned association is no longer valid.

### Interhemispheric gamma synchrony between PV interneurons increases after errors during rule shifts

To examine whether this PV activity, which occurs at critical behavioral timepoints, was associated with long-range gamma synchrony, we used TEMPO (Trans-membrane Electrical Measurements Performed Optically). TEMPO utilizes bulk measurements of fluorescence from the voltage indicator Ace2N-4AA-mNeon (mNeon), to quantify rhythmic activity in specific cell types^28^. We injected AAV into mPFC bilaterally to drive Cre-dependent mNeon expression in *PV-Cre, Ai14* mice, and implanted multimode optical fibers to measure mNeon and tdTomato fluorescence from PV interneurons in the left and right mPFC (Fig. 2a,b; Methods). Some mice were also injected with AAV1-Synapsin-tdTomato (Syn-tdTomato). tdTomato signals from each optical fiber represent non-voltage-dependent reference signals (Fig. 2a,b). We measured voltage signals from PV interneurons in both prefrontal cortices (Fig. 2c) while mice learned an initial association, then learned a new association during a rule shift (Fig. 2d).

**Fig. 2:**
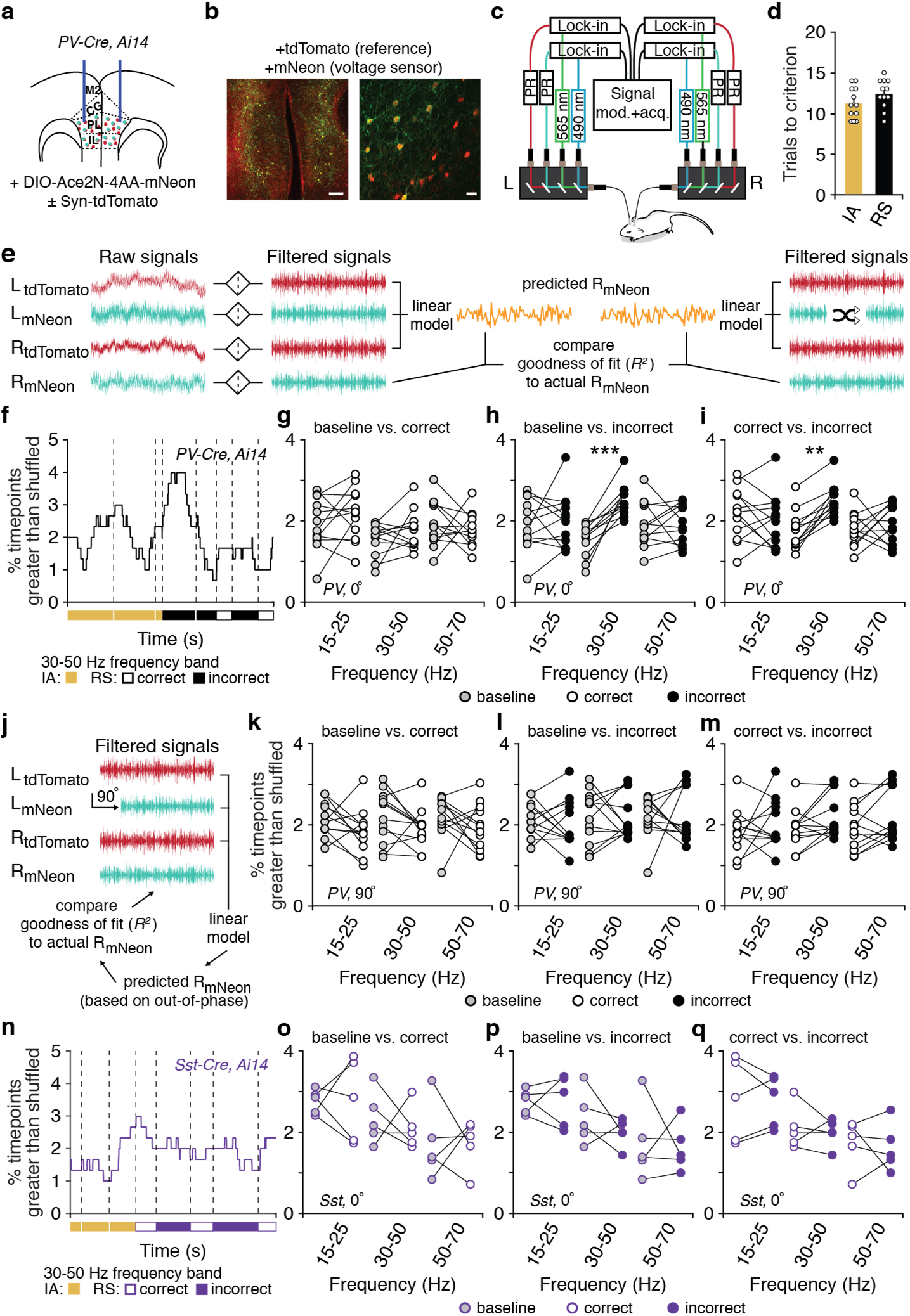
Interhemispheric gamma synchrony of PV interneurons increases after errors during rule shifts. **a**, *PV-Cre, Ai14* mice (*n* = 12) had bilateral AAV-DIO-Ace2N-4AA-mNeon ± AAV-Syn-tdTomato injections and fiber-optic implants in mPFC. **b**, Examples of tdTomato (red) and mNeon (green) fluorescence in a coronal section of mPFC (left), alongside a high power image (right). Scale bars: 100 μm and 25 μm, respectively. **c**, Schematic for dual-site TEMPO imaging. Each fiber-optic implant, for delivering illumination and collecting fluorescence, connects to a mini-cube coupled to two LEDs and two photoreceivers (PR) to separately excite and collect emitted fluorescence from mNeon and tdTomato. Two lock-in amplifiers modulate LED output and demodulate PR signals, which are then acquired by a multichannel real-time signal processor. **d**, Initial association (IA) and rule shift (RS) performance in this cohort. **e**, Overview of dual-site TEMPO analysis: tdTomato and mNeon fluorescence signals from each hemisphere are filtered around a frequency of interest, then both tdTomato signals and one mNeon signal are used to model the second mNeon signal. Performance is compared to models based on shuffled versions of the first mNeon signal. **f**, *R^2^* values, measuring zero-phase lag ~40 Hz interhemispheric PV interneuron synchrony, during the last 3 IA trials and the first 5 RS trials in one mouse. **g**, Synchrony was not different after correct decisions vs. during the baseline period (two-way ANOVA; frequency X condition interaction: *F*_2,44_ = 0.88, *P* = 0.42). **h-i**, 30-50 Hz synchronization was specifically higher after RS errors than during the baseline period (twoway ANOVA; main effect of condition: *F*_1,22_ = 5.2, \**P* = 0.033; frequency X condition interaction: *F*_2,44_ = 7.89, \*\**P* = 0.001; 15-25 Hz: post hoc *t_(66)_* = 0.37, *P* > 0.99; 30-50 Hz: post hoc *t_(66)_* = 4.43, \*\*\**P* = 0.0001; 50-70 Hz: post hoc *t_(66)_* = 0.006, *P* > 0.99) or after RS correct decisions (two-way ANOVA; frequency X condition interaction: *F_2,44_* = 5.36, \*\**P* = 0.008; post hoc *t_(66)_* = 3.53, \*\**P* = 0.002). **j**, Schematic: analysis to measure out-of-phase synchrony. In this case, one mNeon is signal is shifted 90 degrees out-of-phase relative to the other signals, before following the procedure outlined in panel (**e**). **k-m**, Out-of-phase 30-50 Hz synchrony did not differ between the baseline period and RS correct trials (two-way ANOVA; frequency X condition interaction: *F*_2,44_ = 0.023, *P* = 0.98; post hoc *t_(66)_* = 1.31, *P* = 0.59), baseline period and RS incorrect trials (two-way ANOVA; frequency X condition interaction: *F*_2,44_ = 0.11, *P* = 0.90; post hoc *t_(66)_* = 0.049, *P* > 0.99), or correct and incorrect trials (two-way ANOVA; frequency X condition interaction: *F*_2,44_ = 0.075, *P* = 0.93; post hoc *t_(66)_* = 1.17, *P* = 0.74). n, *R^2^* values, measuring zero-phase lag ~40 Hz interhemispheric Sst interneuron synchrony during the last 3 IA trials and the first 5 RS trials in one mouse. **o-q**, Interhemispheric Sst synchrony (*n* = 5) was not different between the baseline period and RS correct trials (two-way ANOVA; main effect of condition: *F*_1,12_ = 0.07, *P* = 0.79; main effect of frequency: *F*_2,12_ = 6.07, \**P* = 0.015; frequency X condition interaction *F*_2,12_ = 0.10, *P* = 0.90), baseline period and RS incorrect trials (two-way ANOVA; main effect of condition: *F*_2,12_ = 0.47, *P* = 0.51; main effect of frequency: *F*_1,12_ = 6.07, \**P* = 0.015; frequency X condition interaction: *F*_1,12_ = 0.26, *P* = 0.78), nor correct and incorrect trials (two-way ANOVA; main effect of condition: *F*_1,12_ = 0.25, *P* = 0.63; main effect of frequency: *F*_2,12_ = 4.66, \**P* = 0.032; frequency X condition interaction: *F*_2,12_ = 0.034, *P* = 0.97). Data are shown as means (**d**); error bars (**d**) denote s.e.m. Two-way ANOVA followed by Bonferroni post hoc comparisons were used. Comparisons were not significant unless otherwise noted.

Fluorescent signals from genetically encoded voltage indicators such as mNeon are contaminated by hemodynamic signals, making their effective signal-to-noise relatively low^28,29^. What this means is that conventional spectral analyses of these signals, e.g., measuring their power or coherence at different frequencies, will be dominated by non-neuronal artifacts. Previous studies have dealt with this issue by using unsupervised methods to separate signal from noise in recordings from a single site^28,29^. However, this approach does not clearly resolve gamma-frequency signals in freely-behaving mice. Here, we were able to overcome this barrier by leveraging dual-site recordings to develop a new method for quantifying synchronization at specific frequencies. Specifically, to quantify zero-phase lag interhemispheric synchronization between left and right mNeon signals, we filtered all signals around a frequency of interest, then used the left mNeon signal, left tdTomato signal, and right tdTomato signal as inputs to a linear model which we used to predict the right mNeon signal (Fig. 2e). At every point in time, we compared the performance of this model to that of 100 models which used the actual tdTomato signals along with time-shifted versions of the left mNeon signal (Fig. 2e). The left and right tdTomato signals should capture the shared sources of noise, e.g., hemodynamic signals, movement artifacts, fiber bending, etc., and the shuffled left mNeon signal should match the degrees of freedom. Thus, the degree to which the model based on the actual left mNeon signal outperforms those based on shuffled left mNeon signal should reflect the degree to which the left mNeon signal carries zero-phase lag information about the right mNeon signal, i.e., the degree of (zero-phase lag) interhemispheric synchronization (at a frequency of interest) between prefrontal PV interneurons. Using this method, we analyzed the first five rule-shift trials in *PV-Cre* mice (Fig. 2f–m). Interhemispheric PV interneuron synchronization in the 30-50 Hz band was significantly higher following errors than during the baseline period or following correct decisions (Fig. 2f–i). Notably, this error-related increase in synchrony was frequency-specific: for both lower (15-25 Hz) and higher (50-70 Hz) frequency bands, interhemispheric PV interneuron synchronization was similar across the baseline period, after correct decisions, and after errors.

Synchronization at an arbitrary phase lag can be expressed as the sum of in-phase and 90 degree out-of-phase components. Therefore, to explore potential synchrony at non-zero phase lags, we also measured synchronization using mNeon signals from PV interneurons in the left mPFC that had been phase-shifted 90 degrees (Fig. 2j–m). In this case, there were no differences in 30-50 Hz synchronization between the baseline period and the periods following correct or incorrect decisions (Figures 2k–m). This indicates that it is specifically the zero-phase lag component of gamma-frequency synchronization that is recruited following rule-shift errors. To confirm that this increase in zero-phase lag synchronization is cell-type specific, we also used TEMPO to measure the voltage dynamics of somatostatin (Sst) interneurons in *Sst-Cre* mice (Fig. 2n–q; Supplementary Fig. 2). Interhemispheric synchrony between prefrontal Sst interneurons was not higher following errors, compared to correct trials or baseline periods. Thus, the synchronization we observed between left and right TEMPO signals is specific for correct vs. incorrect trial outcome, frequency band, phase lag, and cell type — all of which represent controls for potential nonspecific artifacts related to movement, respiration, hemodynamics, etc. The fact that this increase in synchronization was specific for cell type also indicates that it is not a generic feature of mPFC activity; furthermore, the fact that it was specific for frequency band suggests that it is not just indicative of increased PV interneuron activity.

### Increases in gamma synchrony are specific for rule shifts

Next we asked whether this increase in gamma synchrony is specific for the rule-shift aspect of our task, or whether it represents a more generic signal related to error detection or reinforcement. First, we reanalyzed our photometry data, and found that the increases in PV interneuron activity that normally follow rule-shift errors do not occur after errors during learning of an initial association (Fig. 3a–d). Then, we used TEMPO to compare changes in interhemispheric PV interneuron gamma synchrony during learning of an initial association, rule reversal, or rule shift (Fig. 3e–f; Supplementary Fig. 3). Unlike the increase in interhemispheric PV interneuron gamma synchrony we observed following rule-shift errors, there was no difference between PV interneuron gamma synchrony on correct vs. incorrect trials during an initial association (Fig. 3g) or rule reversal (Fig. 3h). Furthermore, 30-50 Hz synchrony was significantly higher on errors during rule shifts than on errors during learning of initial associations (Fig. 3i) or rule reversals (Fig. 3j). Notably, this difference was specific for the 30-50 Hz band and was not present for 15-25 or for 50-70 Hz synchrony.

**Fig. 3:**
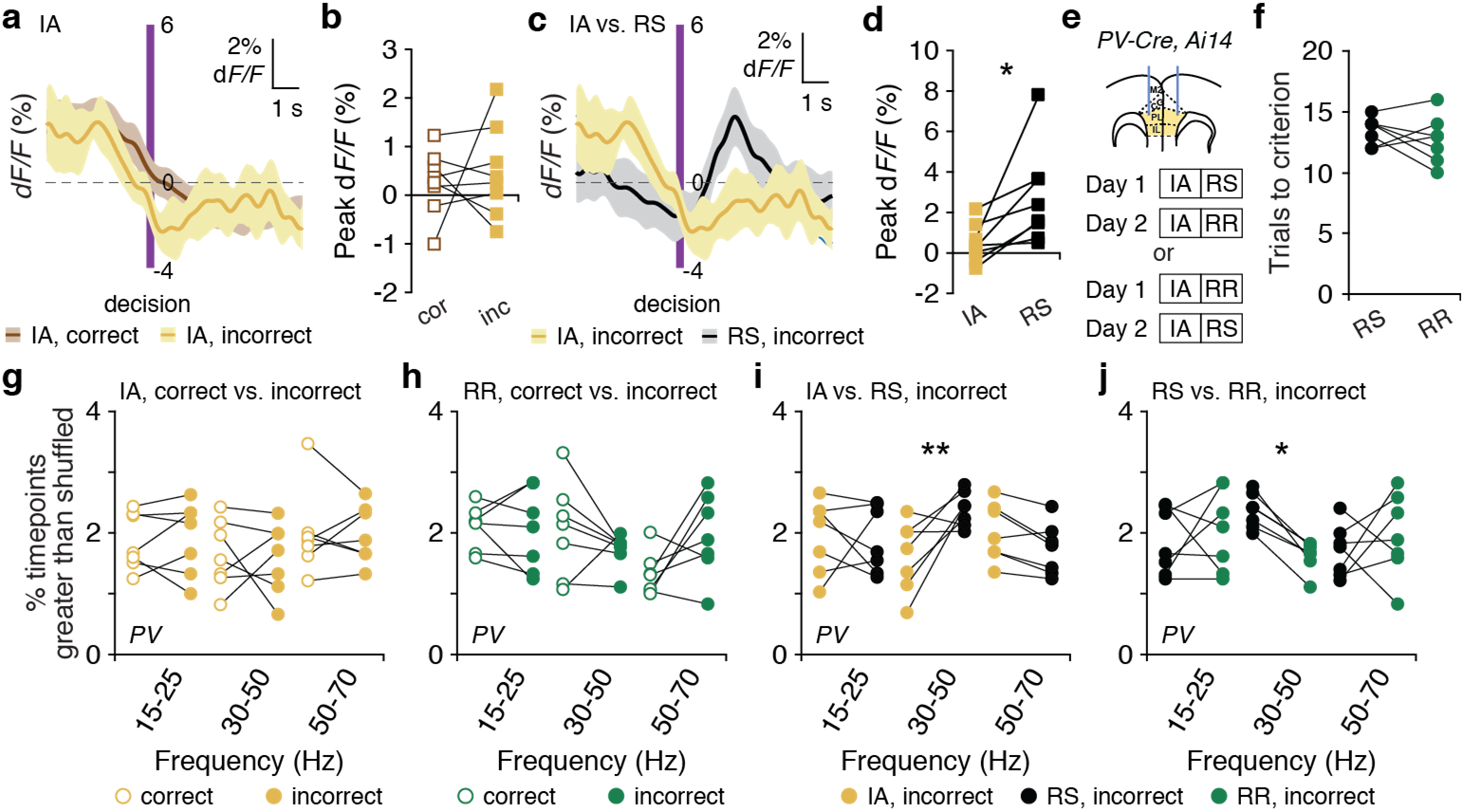
Interhemispheric synchrony does not increase during initial associations and rule reversals. **a**, Averaged PV interneuron photometry signal (*dF/F*), aligned to the time of dig, which indicates a decision, for correct (brown line) and incorrect trials (yellow line) during the initial association (IA) (*n* = 8). **b**, Peak *dF/F* during the 4 sec following the decision. Signals are comparable during correct and incorrect IA trials (two-tailed, paired **t*-test; t_(7)_* = 0.44, *P* = 0.67). **c**, Averaged PV interneuron photometry signal (*dF/F*), aligned to the time of dig, which indicates a decision, for incorrect trials during the IA (yellow line) or rule shift (RS) (black line; *n* = 8). **d**, Peak *dF/F* during the 4 sec following the decision. Signals on incorrect trials are significantly lower during the IA than the RS (two-tailed, paired *t*-test; *t_(7)_* = 2.87, \**P* = 0.024). **e**, *PV-Cre, Ai14* mice (*n* = 7) had bilateral injections of AAV-DIO-Ace2N-4AA-mNeon ± AAV-Syn-tdTomato in mPFC and fiber-optic implants in mPFC. Experimental design: Day 1: IA followed by RS or RR; Day 2: IA followed by the task not performed on Day 1. **f**, *PV-Cre, Ai14* mice performed rule shifts (RS) and rule reversals (RR) in a similar number of trials (twotailed, paired *t*-test; *t_(6)_* = 0.92, *P* = 0.39). **g**, During the IA, synchrony did not differ following correct vs. incorrect trials (two-way ANOVA; main effect of condition: *F*_1,12_ = 0.0002, *P* = 0.99; frequency X condition interaction: *F*_2,24_ = 0.048, *P* = 0.95). **h**, During the RR, synchrony did not differ following correct vs. incorrect trials (two-way ANOVA; main effect of condition: *F*_1,18_ = 0.10, *P* = 0.76; interaction *F*_2,18_ = 4.55, \**P* = 0.025; 15-25 Hz: post hoc *t_(18)_* = 0.25, *P* > 0.99; 30-50 Hz: post hoc *t_(18)_* = 1.71, *P* = 0.32; 50-70 Hz: post hoc *t_(18)_* = 2.50, *P* = 0.07). **i**, Following errors, synchrony was specifically higher for the 30-50 Hz band during RS than IA (two-way ANOVA; condition X frequency interaction: *F*_2,18_ = 6.02, \*\**P* = 0.0099; 30-50 Hz post hoc *t_(18)_* = 3.42, \*\**P* = 0.009). **j**, Following errors, synchrony was specifically higher in the 30-50 Hz band RS than RR (two-way ANOVA; condition X frequency interaction *F_2,18_* = 3.96, \**P* = 0.038; 30-50 Hz post hoc *t_(18)_* = 2.64, \**P* = 0.0499). Data are shown as means (a, c); shading (a, c) denotes s.e.m. Two-way ANOVA followed by Bonferroni post hoc comparisons, unless otherwise noted.

Rule changes (i.e., rule shifts or rule reversals) are uncued. Therefore, the nature of the rule change, and any associated differences in synchrony, should only become apparent over time. To confirm that this is the case, we performed a trial-by-trial analysis of differences in gamma synchrony on incorrect vs. correct trials, during rule shifts vs. rule reversals. Indeed, we found that the tendency for gamma synchrony to be higher on incorrect vs. correct trials during rule shifts but not rule reversals was difficult to discern during the first two trials of a rule change. Then, this difference became significantly larger over the next three trials (Supplementary Fig. 3k).

### Perturbing gamma synchrony disrupts learning during rule shifts

To directly test the functional significance of these increases in synchronization that normally follow rule-shift errors, we would like to be able to artificially manipulate the gamma-frequency synchronization of PV interneurons in the left vs. right mPFC. We previously showed that delivering 40 Hz optogenetic stimulation to interneurons *in-phase* across the left and right mPFC does not impair the ability of normal mice to learn a new association during a rule shift^14^. Critically, this optogenetic stimulation was relatively weak. Whereas other studies have examined behavioral effects of 40 Hz stimulation of PV interneurons using light powers of ~5-7 mW^22,24^, our study used ~1 mW. Indeed, one other study that also stimulated PV interneurons at 40 Hz using ~1 mW observed only slight changes in excitatory neuron firing rates^23^; similarly, we found that outside of a small bump at 40 Hz, the LFP power spectrum was minimally altered by 1 mW stimulation of PV interneurons (Supplementary Fig. 4). Thus, one strategy would be to use relatively weak optogenetic stimulation to entrain PV interneurons in the left vs. right mPFC with different phase relationships, to determine whether these patterns, which should elicit similar changes in *levels* of inhibition but differentially affect interhemispheric synchronization, elicit similar vs. distinct effects on behavior.

Based on this logic, we performed two experiments. First, we found that stimulating prefrontal PV interneurons at 40 Hz but *out-of-phase* across the two hemispheres disrupts learning of new associations during rule shifts (Fig. 4a–e). Mice receiving out-of-phase interhemispheric stimulation took significantly longer to learn compared to either control (eYFP-expressing) mice, or to themselves on a different day when no stimulation was delivered. We also characterized the types of errors made during learning. Because odor-texture pairings vary randomly from trial to trial, during rule shifts mice can make both *perseverative* and *random* errors. Perseverative errors occur when the originally-rewarded cue and the newly-rewarded cue are located in different bowls, and the mouse chooses the originally-rewarded cue. In contrast, random errors occur when the originally-rewarded cue and the newly-rewarded cue are located in the same bowl, but the mouse chooses the other bowl. Out-of-phase 40 Hz stimulation caused mice to make significantly more perseverative errors, again compared to either the no stimulation condition or control mice (Fig. 4b,c; Supplementary Fig. 5). Next, in a different cohort of *PV-Cre* mice, we compared the effects of a) out-of-phase 20 Hz stimulation, b) out-of-phase 40 Hz stimulation, and c) in-phase 40 Hz stimulation, delivered to PV interneurons in the left vs. right mPFC during a rule shift (Fig. 4d). Out-of-phase 20 Hz stimulation had no effect on learning of a new association during the rule shift, i.e., mice learned normally, in 10-15 trials (Fig. 4e; compare to Fig. 2d). Again, out-of-phase 40 Hz stimulation markedly disrupted learning during the rule shift and increased perseveration, but when we delivered inphase 40 Hz stimulation on the following day, the same mice once again learned normally (Fig. 4e; Supplementary Fig. 5).

**Fig. 4.**
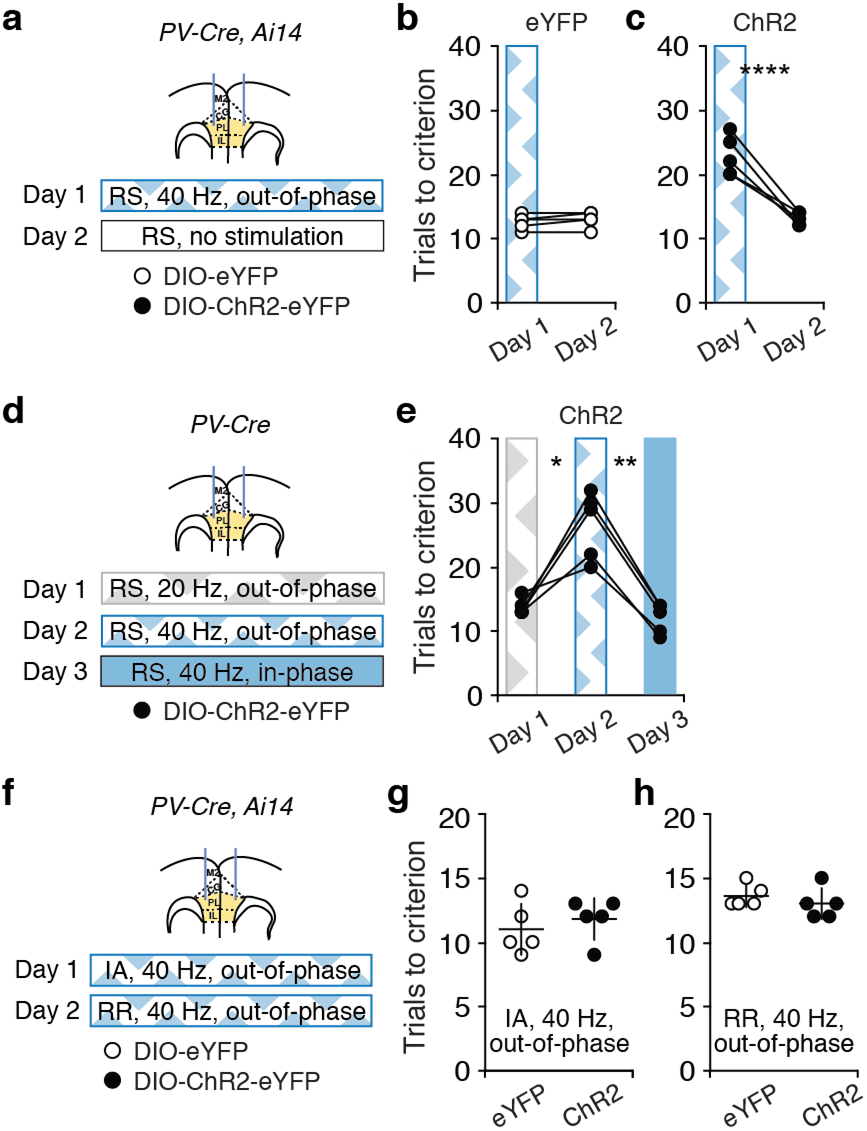
Out-of-phase, but not in-phase, gamma-frequency stimulation of PV interneurons disrupts learning during rule shifts but not during initial associations or rule reversals. **a**, *PV-Cre, Ai14* mice had injections of AAV-DIO-eYFP (*n* = 5) or AAV-DIO-ChR2-eYFP (*n* = 5) in mPFC and fiber-optic implants bilaterally in mPFC. Experimental design: Day 1: out-of-phase 40 Hz stimulation during the rule shift (RS); Day 2: no stimulation. **b-c**, Out of phase 40 Hz stimulation impairs rule-shifting in ChR2-expressing mice compared to eYFP-expressing controls (two-way ANOVA; main effect of day: *F*_1,8_ = 40.2, \*\*\**P* = 0.0002; main effect of virus: *F*_1,8_ = 32.5, \*\*\**P* = 0.0005; day X virus interaction: *F*_1,8_ = 47.3, \*\*\**P* = 0.0001). (**b**) Performance of eYFP-expressing controls did not change from Day 1 to 2 (post hoc *t_(8)_* = 0.38, *P* > 0.99). (**c**) Out-of-phase 40 Hz stimulation of PV interneurons across hemispheres during the RS on Day 1 impaired rule shifts in ChR2-expressing mice, compared to no stimulation on Day 2 (post hoc *t_(8)_* = 9.34, ****P < 0.0001). **d**, *PV-Cre* mice (*n* = 5) had bilateral injections of AAV-DIO-ChR2-eYFP and fiber-optic implants in mPFC. Experimental design: Day 1: out-of-phase 20 Hz stimulation; Day 2: out-of-phase 40 Hz stimulation; Day 3: in-phase 40 Hz stimulation. **e**, Out-of-phase 40 Hz stimulation (Day 2) impairs rule shifts relative to out-of-phase 20 Hz stimulation (Day 1) or in-phase 40 Hz stimulation (Day 3) (one-way repeated measures ANOVA followed by Tukey’s multiple comparisons test, main effect of treatment: *F*_1.005,4.021_ = 33.5, \*\**P* = 0.004; Day 1 vs. Day 2: \**P* = 0.021, Day 2 vs. Day 3: \*\**P* = 0.001, Day 1 vs. Day 3: *P* = 0.45). **f**, *PV-Cre, Ai14* mice had bilateral injections of AAV-DIO-eYFP (*n* = 5) or AAV-DIO-ChR2-eYFP (*n* = 5) and fiber-optic implants in mPFC. Experimental design: Day 1: out-of-phase 40 Hz stimulation during the initial association (IA); Day 2: no stimulation during the IA, followed by out-of-phase 40 Hz stimulation during the rule reversal (RR). **g**, Out-of-phase 40 Hz stimulation does not affect the ability of ChR2-expressing mice to learn an IA (twotailed, unpaired *t*-test; compared to control eYFP-expressing mice; *t_(8)_* = 0.69, *P* = 0.51). **h**, Out-of-phase 40 Hz stimulation does not affect the ability of ChR2-expressing mice to learn a RR (twotailed, unpaired *t*-test; compared to control eYFP-expressing mice; *t_(8)_* = 0.89, *P* = 0.40). Data are shown as means (**g-h**); error bars (**g, h**) denote s.e.m. Two-way ANOVA followed by Bonferroni post hoc comparisons, unless otherwise noted.

Thus, the ability to learn new associations during rule shifts depends critically on the phase relationship of gamma-frequency activity between PV interneurons in the left and right mPFC. Delivering stimulation to perturb the zero-phase lag 30-50 Hz interhemispheric synchrony that normally occurs after rule-shift errors disrupts learning during that rule shift. By contrast, our previous work and new experiments both confirmed that when the same pattern of stimulation is delivered in-phase, rather than out-of-phase across hemispheres, learning during a rule shift is normal. Disrupting synchronization at lower frequencies also has no effect, suggesting that synchronization specifically within the gamma band is important for this form of learning.

### Perturbing gamma synchrony does not affect initial associations or rule reversals

Given that we observed increased gamma synchrony during rule shifts, but not during initial associations or rule reversals, we wondered whether gamma synchrony is specifically necessary for learning during rule shifts, or whether perturbing gamma synchrony would also disrupt the acquisition of these other types of associations. To examine this, we stimulated prefrontal PV interneurons at 40 Hz, but out-of-phase across the two hemispheres during an initial association or rule reversal. In contrast to its ability to disrupt learning during rule shifts, out-of-phase 40 Hz stimulation did not affect learning during initial associations or rule reversals (Fig. 4f—h). Together with our earlier TEMPO experiments, this shows that interhemispheric gamma synchrony is specifically recruited by and necessary for learning of new associations during rule shifts. Unlike initial associations or rule reversals, rule shifts require mice to stop utilizing one set of cues, and to instead learn a new association which reappraises the behavioral significance of cues that were previously irrelevant to the outcome of each trial.

### Gamma synchrony fails to increase in mutant mice that exhibit perseveration

Finally, to evaluate how changes in gamma-frequency synchrony might contribute to, or be targeted to alleviate, pathological phenotypes, we exploited *Dlx5/6^+/-^* mutant mice, which have abnormal PV interneuron physiology, deficient task-evoked gamma oscillations, and perseverative behavior during rule shifts^14^. Using fiber photometry (Fig. 5a–f; Supplementary Fig. 6), we found that the prominent error-related activity we observed in wild-type (*Dlx5/6^+/+^*) PV interneurons during rule shifts was significantly attenuated in *Dlx5/6* mutants (Fig. 5e,f). By contrast, PV interneuron activity following correct decisions did not differ between wild-type mice and *Dlx5/6* mutants (Fig. 5c,d). Whereas dual-site TEMPO imaging from PV interneurons in wild-type mice had revealed an increase in 30-50 Hz interhemispheric synchronization on incorrect trials relative to the baseline period or correct trials, synchronization in mutant mice was not higher after rule-shift errors vs. correct decisions (Fig. 5g,h; Supplementary Fig. 7). Correspondingly, the normal increase in 30-50 Hz interhemispheric PV interneuron synchronization following rule-shift errors (relative to correct trials) was significantly attenuated in *Dlx5/6^+/-^* mice compared to wild-types (Fig. 5j). Notably, this difference was specific for frequency band, task, and cell type.

**Figure 5:**
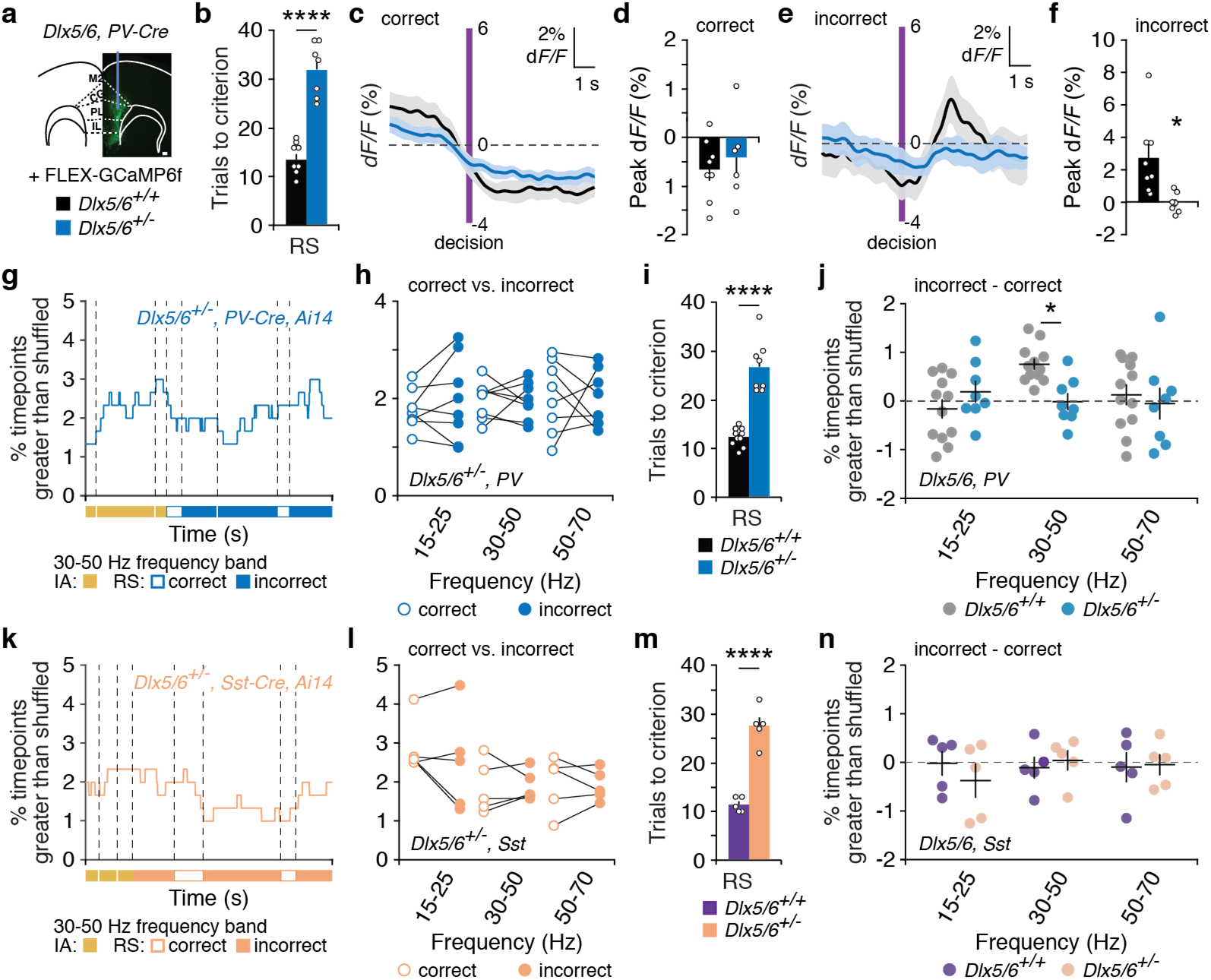
Interhemispheric gamma synchrony fails to increase during rule shifts in mutant mice. **a**, FLEX-GCaMP6f expression in a *Dlx5/6^+/-^, PV-Cre* mouse (scale bar, 100 μm). **b**, Rule-shifting (RS) performance is impaired in mutant mice (blue; *n* = 7) compared wild-type (*Dlx5/6^+/+^*) littermates (black; *n* = 8; two-tailed, unpaired *t*-test; *t_(13)_* = 7.82, *\*\*\*\*P* < 0.0001). **c**, Averaged *dF/F* from PV interneurons in mutant (blue) vs. wild-type (black) mice, aligned to the time of correct decisions. **d**, Peak PV interneuron *dF/F* values during the 4 sec following correct decisions during rule shifts were similar in *Dlx5/6^+/-^* (blue) vs. wild-type (black) mice (two-tailed, unpaired *t*-test; *t_(13)_* = 0.66, *P* = 0.52). **e**, Averaged *dF/F* from PV interneurons in mutant (blue) vs. wild-type (black) mice, aligned to the time of incorrect decisions. **f**, Peak dF/F from PV interneurons during the 4 sec following incorrect decisions is significantly decreased in *Dlx5/6^+/-^* mice (blue) compared to wild-types (black) (two-tailed, unpaired *t*-test; *t_(8.085)_* = 3.18, \**P* = 0.01). **g**, *R^2^* values, measuring zero-phase lag ~40 Hz interhemispheric interneuron synchronization between TEMPO signals from PV interneurons in mutant mice during the last 3 IA and first 5 RS trials in one *Dlx5/6^+/-^* mouse. **h**, In *Dlx5/6^+/-^* mice, interhemispheric PV interneuron synchronization was not different following errors vs. correct decisions (two-way ANOVA; main effect of condition: *F*_1,21_ = 0.09, *P* = 0.77; condition X frequency interaction: *F*_2,21_ = 0.29, *P* = 0.75). **i**, Rule-shift performance is impaired in mutant mice (blue; *n* = 8) compared to wild-type (black) littermates (*n* = 12; two-tailed, unpaired *t*-test; *t_(8.071)_* = 7.40, \*\*\*\**P* < 0.0001). **j**, Increases in PV interneuron synchrony following errors (relative to synchrony after correct decisions) are significantly attenuated in mutants compared to wild-type littermates, specifically in the 30-50 Hz frequency band (two-way ANOVA; genotype X frequency interaction: *F*_2,54_ = 3.74, \**P* = 0.030; 15-25 Hz: post hoc *t_(54)_* = 1.21, *P* = 0.69; 30-50 Hz: post hoc *t_(54)_* = 2.65, \**P* = 0.032; 50-70 Hz: *t_(54)_* = 0.62, *P* > 0.99). **k**, *R^2^* values, measuring zero-phase lag ~40 Hz interhemispheric Sst interneuron synchronization during the last 3 IA and first 5 RS trials in one *Dlx5/6^+/-^* mouse. **l**, In mutants (*n* = 5), interhemispheric Sst interneuron synchrony is similar following correct vs. incorrect decisions (two-way ANOVA; main effect of condition: *F*_1,12_ = 0.25, *P* = 0.63; main effect of frequency: *F2,12* = 4.6, \**P* = 0.0318; condition X frequency interaction: *F*_2,12_ = 0.034, *P* = 0.97). **m**, Rule-shifting is impaired in mutants (*n* = 5) compared to wild-type littermates (*n* = 5; two-tailed, unpaired *t*-test; *t_(8)_* = 8.64, *\*\*\*\*P* < 0.0001). **n**, Changes in Sst interneuron synchrony following errors (relative to synchrony after correct decisions) are not different in mutant vs. wild-type mice (two-way ANOVA; main effect of genotype: *F*_1,16_ = 0.06, *P* = 0.82; genotype X frequency interaction: *F*_2,16_ = 0.54, *P* = 0.59). Data are shown as means (**b–f, i–j, m–n**); error bars (**b, d, f, i–j, m–n**) and shading (**c, e**) denote s.e.m. Two-way ANOVA followed by Bonferroni post hoc comparisons were used, unless otherwise noted.

Increased synchrony after rule-shift errors (relative to correct trials) was deficient for the 30-50 Hz band but not for 15-25 or for 50-70 Hz (Fig. 5j). Furthermore, in mutants, synchronization was not different from wild-types during the baseline period or during learning of an initial association (Supplementary Fig. 8). Lastly, in *Dlx5/6* mutants, deficits in gamma-frequency synchronization occurred specifically within PV interneurons. Using dual-site TEMPO to measure interhemispheric synchronization of Sst interneurons, we did not observe differences between wild-type and *Dlx5/6* mutant mice (Fig. 5k–n; Supplementary Fig. 6c,d; Supplementary Fig. 7).

### Only in-phase gamma-frequency PV interneuron stimulation rescues perseveration

We previously found that 40 Hz stimulation of mPFC interneurons rescues learning during rule shifts in *Dlx5/6* mutant mice. Importantly, in those experiments, delivering the same amount of stimulation using frequencies above (125 Hz) and below (12.5 Hz) the gamma band was ineffective^14^. Now, our new results suggest that increasing gamma *synchrony* between prefrontal PV interneurons during the early portions of a rule shift may be particularly critical for stimulation to facilitate learning in mutant mice. We tested this in three ways. First, we found that restricting optogenetic stimulation to only PV interneurons and just the initial portion of the rule shift (the first five trials), was still sufficient to rescue learning during rule shifts in *Dlx5/6* mutants (Fig. 6a–c; Supplementary Fig. 6). Second, we tested the effects of stimulating PV interneurons with the same frequencies and phase relationships tested earlier in wild-type (*Dlx5/6^+/+^*) mice. 20 or 40 Hz stimulation that was out-of-phase across the two hemispheres did not improve learning during rule shifts in mutant mice (Fig. 6d–f; Supplementary Fig. 6). However, consistent with our earlier findings^14^, in-phase 40 Hz stimulation rescued learning during rule shifts in *Dlx5/6* mutants (Fig. 6f; Supplementary Fig. 6). (Importantly, we previously showed that in the absence of optogenetic stimulation, learning during rule shifts in *Dlx5/6* mutant mice does not improve over three consecutive days of testing^14^). These results show that gamma-frequency activity in prefrontal PV interneurons is not sufficient to facilitate learning during rule shifts unless it is precisely synchronized across hemispheres. Finally, to determine the cell-type specificity of this behavioral effect, we delivered in-phase, 40 Hz stimulation bilaterally to *Dlx5/6^+/-^*, *Sst-Cre* mice and found that this did not improve learning during rule shifts (Supplementary Fig. 9).

**Fig. 6:**
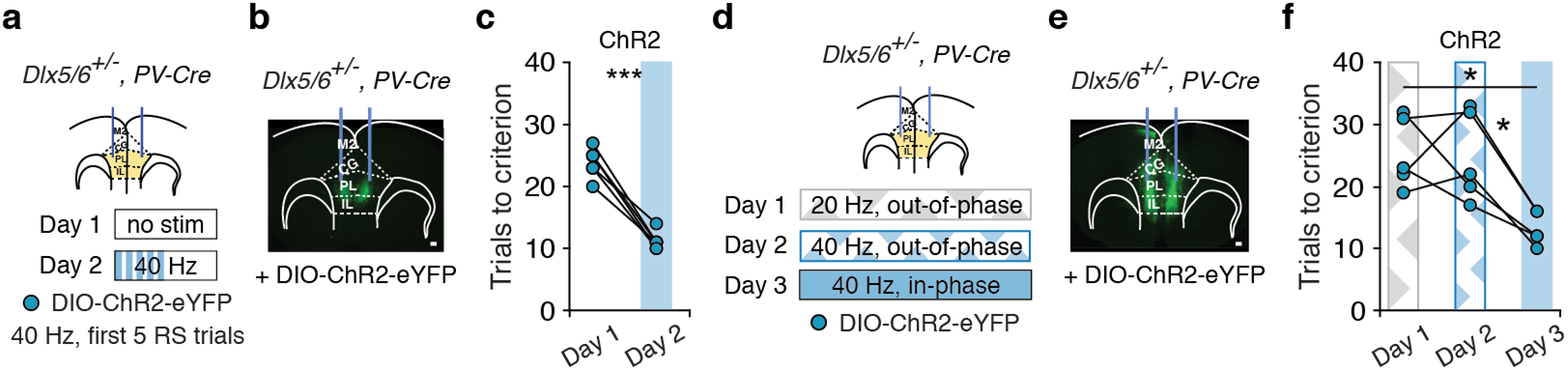
Restoring interhemispheric PV interneuron gamma synchrony is required to rescue rule shifting in *Dlx5/6^+/-^* mutant mice. **a**, *Dlx5/6^+/-^, PV-Cre* mice (*n* = 6) had bilateral AAV-DIO-ChR2-eYFP injections and fiber-optic implants in mPFC. Experimental design: Day 1: no stimulation; Day 2: in-phase 40 Hz stimulation during the first 5 RS trials. **b**, ChR2-eYFP expression in the mPFC of a *Dlx5/6^+/-^, PV-Cre* mouse (scale bar, 100 μm). **c**, In-phase 40 Hz stimulation on Day 2 normalizes rule-shifting in mutant mice (two-tailed, paired *t*-test; *t_(5)_* = 10.3, \*\*\**P* = 0.0001). **d**, *Dlx5/6^+/-^*, *PV-Cre* mice (*n* = 5) had bilateral AAV-DIO-ChR2-eYFP injections and fiber-optic implants in mPFC. Experimental design: Day 1: out-of-phase 20 Hz stimulation; Day 2: out-of-phase 40 Hz stimulation; Day 3: in-phase 40 Hz stimulation. **e**, ChR2-eYFP in the mPFC of a *Dlx5/6^+/-^, PV-Cre* mouse (scale bar, 100 μm). **f**, In mutant mice, in-phase 40 Hz stimulation (Day 3), but not out-of-phase 40 Hz stimulation (Day 2), rescues rule-shifting performance (one-way repeated measures ANOVA followed by Tukey’s multiple comparisons test, main effect of treatment: F_1.291,5.164_ = 10.6, \**P* = 0.019; Day 1 vs. Day 2: *P* = 0.99, Day 2 vs. 3: \**P* = 0.016, Day 1 vs. 3: \**P* = 0.016). Two-way ANOVA followed by Bonferroni post hoc comparisons were used, unless otherwise noted.

### A pharmacological intervention, which rescues learning during rule shifts, specifically increases gamma synchrony

Optogenetic stimulation might be viewed as an ‘artificial’ mechanism for altering gamma synchrony. Therefore we explored whether manipulations which engage endogenous physiological mechanisms might also increase gamma synchrony and elicit similar behavioral effects. For this, we used low – sub-anxiolytic and sub-sedative – doses of the benzodiazepine clonazepam. Our previous study^14^, showed that low-dose clonazepam (0.0625 mg/kg, I.P.), like 40 Hz optogenetic stimulation, normalizes learning during rule shifts in *Dlx5/6^+/-^* mice. Therefore, we now used voltage imaging to assess whether clonazepam increases gamma synchrony during rule shifting (Fig. 7). Indeed, compared to vehicle treatment, clonazepam increased interhemispheric gamma synchrony between prefrontal PV interneurons in *Dlx5/6^+/-^* mice. Moreover, this occurred specifically after errors during rule shifts; there was no increase in synchrony during the baseline period or after correct decisions. Increased synchrony was also specific for the gamma frequency band.

**Fig. 7:**
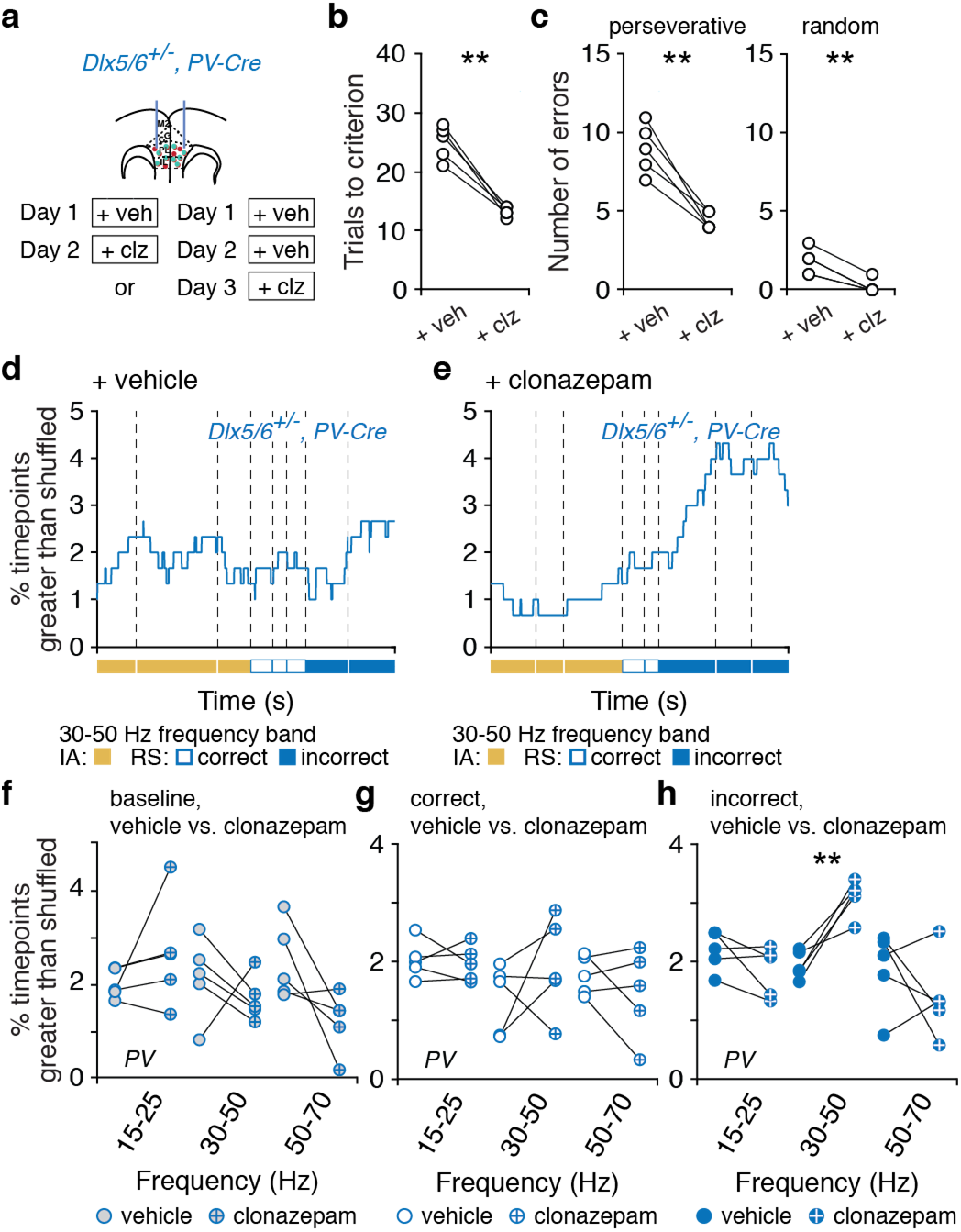
Low-dose clonazepam increases interhemispheric gamma synchrony during rule shifts in mutant mice. **a**, *Dlx5/6^+/-^, PV-Cre* mice (*n* = 5) had bilateral AAV-DIO-Ace2N-4AA-mNeon ± AAV-Syn-tdTomato injections and fiber-optic implants in mPFC. Experimental design: all mice received vehicle only (veh) on Day 1. On Day 2, some mice received clonazepam (clz; *n* = 3); others received veh. On Day 3, we administered clz to those mice which had received veh on Day 2 (*n* = 2). **b**, Low-dose clonazepam normalizes rule-shifting in mutant mice (two-tailed, paired *t*-test; *t_(4)_* = 8.07, \*\**P* = 0.0013). **c**, Low-dose clonazepam decreases perseverative and random errors (two-tailed, paired *t*-test; *t_(4)_* = 6.15, \*\**P* = 0.0036 for perseverative, *t_(4)_* = 6.53, \*\**P* = 0.0028 for random). **d**, *R^2^* values, measuring zero-phase lag ~40 Hz interhemispheric interneuron synchronization between TEMPO signals from PV interneurons in one *Dlx5/6^+/-^* mouse, during the last 3 IA and first 5 RS trials, in the vehicle condition. **e**, *R^2^* values, measuring zero-phase lag ~40 Hz interhemispheric interneuron synchronization between TEMPO signals from PV interneurons in the same mutant mouse, during the last 3 IA and first 5 RS trials in the clonazepam condition. **f**, During the baseline period, synchrony did not differ between the vehicle and clonazepam conditions (two-way ANOVA; main effect of treatment: *F*_1,21_ = 0.0009, *P* = 0.98; frequency X treatment interaction: *F*_2,21_ = 0.86, *P* = 0.44). **g**, Synchrony did not differ following correct trials in the vehicle and clonazepam conditions (two-way ANOVA; main effect of treatment: *F*_1,8_ = 0.09, *P* = 0.77; interaction *F*_2,16_ = 1.48, *P* = 0.26; 15-25 Hz: post hoc *t_(24)_* = 0.18, *P* > 0.99; 30-50 Hz: post hoc *t_(24)_* = 1.55, *P* = 0.41; 50-70 Hz: post hoc *t(24)* = 0.87, *P* > 0.99). **h**, Following RS errors, synchrony was specifically higher in the clonazepam condition for the 30-50 Hz band (two-way ANOVA; treatment X frequency interaction: *F*_2,12_ = 8.63, \*\**P* = 0.0048; 30-50 Hz post hoc *t(12)* = 1.11, \*\**P* = 0.009). Two-way ANOVA followed by Bonferroni post hoc comparisons were used, unless otherwise noted.

## Discussion

It has been suggested that the key function of gamma synchrony is to regulate communication between groups of neurons in order to dynamically reconfigure circuits and generate diverse forms of behavior^7,9,12,13,30^. However, the functional significance of synchronization across neuronal structures, particularly at gamma-frequencies, has long remained one of the most controversial topics in systems neuroscience^31,11,32,18,33,7,8^. Our results directly address this controversy. Specifically, we found a double dissociation between therapeutic or disruptive effects elicited by in-phase vs. out-of-phase stimulation. This confirms that certain aspects of behavior depend on interhemispheric gamma synchrony between PV interneurons, not just rhythmic inhibition in local circuits. Furthermore, in both stimulation and imaging experiments, gamma synchrony was not involved in generic aspects of learning, decision-making, or flexibility, but rather specifically contributed to the formation of new associations based on familiar cues that were previously irrelevant to behavioral outcomes. This study focuses on normal behavior in wild-type mice, but our experiments in *Dlx5/6* mutant mice independently validate (using imaging, optogenetics, and pharmacology) this relationship between gamma synchrony and learning during rule shifts. Notably, this type of learning occurs without extensive prior training, making it very different from the switching between well-learned behaviors that is commonly studied in mice. This ability to develop new strategies which reappraise the salience of external cues is critical for behavioral adaptation to a changing environment.

### A new method for extracting synchrony from genetically encoded voltage indicators

As outlined above, conventional analyses, e.g., power and coherence, cannot be directly applied to signals from genetically encoded voltage indicators, due to their low signal-to-noise. Previous studies have dealt with this issue by using unsupervised approaches, e.g., independent components analysis, to separate signal from noise, but this has not been successful for high-frequency signals (e.g., gamma-frequency signals) in freely-moving mice. In particular, multiple sources of artifacts (fiber bending, movement, changes in blood flow, changes in blood oxygenation, etc.) will differentially contribute to noise at high vs. low frequencies, and the relative amplitudes of these vary over time. As a result, it would be ideal to separate the contributions of signal and noise in a frequency-specific and dynamic manner. Furthermore, in order to estimate synchrony between two signals, e.g., voltage signals from the left and right mPFC, it is not necessary to extract the complete voltage traces, but only the component of the left mPFC voltage trace which is coherent with (i.e., predictive of) the right mPFC voltage. Our new method derives directly from these principles. We filtered mNeon and tdTomato signals within a specific frequency band, and determined how well the left mNeon signal (together with both tdTomato signals) predicts the right mNeon signal within a specific time window. By comparing the strength of prediction obtained using the actual left mNeon signal to that obtained using shuffled versions of that signal, we are able to quantify how much information the left mNeon signal carries about the right mNeon signal. Furthermore, because we included the reference (tdTomato) signals in both cases, we are specifically quantifying information that is present in mNeon signals but not in simultaneously recorded tdTomato signals, i.e., voltage-dependent information.

While currently available genetically encoded voltage indicators are certainly imperfect, several control experiments confirm that we are in fact measuring task-dependent changes in gamma-frequency synchronization between PV interneurons. Specifically, we found that increased synchrony after errors is specific for: 1) the gamma band, 2) rule shifts, and 3) PV interneurons. If the increased synchrony we observed was driven by non-neuronal artifacts, the intrinsic dynamics of mNeon, etc., then increased synchrony should have been present in both PV and Sst interneurons. If increased synchrony was driven by nonspecific aspects of PV neuron activity (as opposed to gamma-frequency activity), we should have seen increased synchrony in other frequency bands. And if this increased synchrony was driven by nonspecific aspects of our task, e.g., generic reward or error signals, movements that mice make after errors, etc., it should have been observed during rule reversals and initial associations as well as during rule shifts. Finally, we only observed increased gamma synchrony between in-phase mNeon signals, and not when one mNeon signal was shifted 90 degrees out-of-phase (~6 msec). This confirms that this increased ability to predict one mNeon signal using the other is due to actual synchrony between the signals, not just the fact that they have similar autocorrelations, second-order statistics, etc., all of which would be the same for the in-phase and 90 degree out-of-phase mNeon signals.

The previous observations all represent negative controls, which confirm the specificity of our method for quantifying gamma synchrony within a particular cell-type. Using this approach, we also found that the benzodiazepine clonazepam enhances gamma synchrony. This represents a positive control, which confirms that our method is sensitive to manipulations which are known to enhance PV interneuron output and gamma oscillations.

Our analytical approach could easily be applied to signals from many other types of genetically encoded voltage indicators. At the same time, this approach complements the sorts of measurements that are possible using electrophysiological recordings. Specifically, genetically encoded voltage indicators are ideal for measuring patterns of activity, which manifest specifically at the mesoscale level and within sparse cell types. This is distinct from examining how individual neurons encode information within the phase of their firing, etc.

### The significance of zero-phase lag synchronization

As noted above, increased gamma synchrony following rule shift errors was present between simultaneously recorded mNeon signals, but not when one signal was shifted 90 degrees out-of-phase. A rhythmic signal *X*, that has an arbitrary phase difference relative to a second signal *Y*, can be expressed as the sum of 0 degree (in phase) and 90 degree out-of-phase components. Thus, if population-level PV interneuron activity in the left and right mPFC had a phase difference significantly different from zero, we should have observed an increase in synchrony even when one mNeon signal was shifted 90 degrees out-of-phase. As can be seen from Fig. 2l,m, this was not the case, indicating that the population-level phase lag is close to zero. This may seem puzzling for two reasons. First, it is commonly assumed that synaptic communication between the hemispheres should be associated with a time delay that produces a phase difference. In fact, it is relatively easy for zero-phase lag synchrony to emerge in bidirectionally coupled oscillators, even when they communicate with a significant delay. Consider the simple example of two phase oscillators, representing the two hemispheres, that emit output (spikes) upon completing each cycle. Assume this output reaches the other oscillator after a quarter-cycle delay (for 40 Hz this is ~6 msec), perturbing the phase of the post-synaptic oscillator by an amount proportional to the cosine of its current phase. (Such coupling is not hard to imagine: suppose that output which arrives when excitatory neuron firing is approaching its peak, recruits more excitation, accelerating the next cycle, whereas output arriving later, when inhibitory neurons have been recruited, mainly increases inhibition, delaying the next cycle). For weak coupling, this system exhibits zero-phase lag synchrony, even though communication between these two oscillators (e.g., the left and right mPFC) occurs with a quarter cycle synaptic delay.

A second potential concern is about the potential function of zero-phase lag synchrony. We have previously shown that inputs which are modulated at gamma frequency transmit greater information to downstream neurons than do non-rhythmic inputs^9^. Suppose that the left and right mPFC converge on a common downstream target, e.g., mediodorsal thalamus or striatum. Then, when they are synchronized with zero-phase lag, inputs from the left and right mPFC hemispheres will summate in downstream neurons in a manner that preserves the gamma-frequency modulation present within each individual signal. By contrast, when gamma-frequency is out-of-phase across the hemispheres, the rhythmic modulation of their summated input will be degraded, compromising the transmission of information to downstream targets. Thus, interhemispheric gamma synchrony may potentiate prefrontal outputs to other regions that serve to update the behavioral salience of external cues.

### The functional impacts of perturbing vs. inducing gamma synchrony

Optogenetic stimulation and inhibition are commonly used to test the causal significance of specific patterns of neural activity. However, any optogenetic manipulation induces firing patterns that are, by definition, ‘artificial.’ In this context, several observations suggest that our optogenetic results are informative about normal circuit function, rather than simply inducing non-physiological states. First, we used relatively weak levels of optogenetic stimulation that we and others have found does not markedly alter overall levels of circuit activity. Second, we delivered exactly the same pattern of stimulation to each PFC, either in- or out-of-phase; only out-of-phase stimulation disrupted behavior in normal mice. Behavior was completely normal during in-phase stimulation. Thus, the behavioral effects of stimulation cannot be attributed to excessive PV interneuron firing, or the hypersynchronous firing of PV interneurons within one hemisphere. Rather, the behavioral effects (disrupted learning during rule shifts) must be due to the induction of artificial (nonzero) phase differences, which would disrupt normally occurring patterns of zero-phase lag synchrony.

Finally, in-phase stimulation, which does not affect behavior in normal mice, does rescue learning during rule shifts in mutant (*Dlx5/6^+/-^*) mice. This same behavioral effect can be produced using very low (sub-anxiolytic and sub-sedative) doses of the benzodiazepine clonazepam^14^. Here, we found that low-dose clonazepam restores the increase in gamma synchrony that is normally seen in wild-type mice after errors during rule shifts. This suggests that in-phase stimulation is functionally similar to clonazepam, which acts by enhancing endogenous PV interneuron output. This supports the idea that this form of optogenetic stimulation reproduces physiologically and therapeutically-relevant states, rather than creating aberrant ones. Thus, even though no optogenetic experiment perfectly recapitulates normal patterns of firing, the results of optogenetic experiments can be informative about how specific aspects of normally-occurring activity (in this case, zero-phase lag interhemispheric gamma synchrony between PV interneurons) contribute to behavior.

### Effects of gamma synchrony are specific for PV vs. SST interneurons

Synchronized gamma-frequency activity has long been attributed to PV interneurons^9,34-36^. However, recent studies have shown that in visual cortex, Sst interneurons are responsible for synchronized rhythmic activity at low ends of the gamma-frequency range^37,38^. Our voltage imaging and optogenetic experiments did not find evidence that synchronized gamma-frequency activity in prefrontal Sst interneurons contributes to learning during rule shifts. This is broadly consistent with work suggesting that PV and Sst interneurons play fundamentally different roles in sensory circuits^39–41^. This may reflect the fact that PV and Sst interneurons generate functionally-distinct forms of circuit inhibition, possibly by innervating different classes of pyramidal neurons^42^ as well as different subcellular domains.

### Relevance to disease

Deficits in PV interneurons and gamma oscillations are key pathophysiological features of schizophrenia^43–45^, which has at its core, deficits in prefrontal-dependent cognitive domains^46,47^.

Consistent with the hypothesis that the deficits in PV interneurons and gamma synchrony contribute to these cognitive deficits, we found that deficits in interhemispheric gamma synchrony are a key driver of cognitive deficits in mutant mice, which model key aspects of schizophrenia. Deficits in PV interneurons and gamma synchrony may also contribute to cognitive deficits in Alzheimer’s disease^48^ and recent studies suggest that driving synchronized gamma oscillations may ameliorate both behavioral and neuropathological aspects of this disorder^49,50^. Our findings suggest that interventions which restore gamma oscillations may treat disease-related cognitive deficits, but only when they involve the proper cell types and reproduce endogenous patterns of long-range synchronization.

In individuals at high risk for psychosis, deficits in the ability to learn new associations based on previously irrelevant cues (measured by the WCST) are strongly correlated with impairments in *insight*, the capacity to appraise and modify distorted beliefs about anomalous experiences^51^. Impaired insight is believed to play a central role in the development and maintenance of psychosis^52^. This suggests that gamma synchrony may be relevant to understanding psychosis itself, as well as cognitive dysfunction, in schizophrenia and related disorders.

### Data and code availability

The data and code that support the findings of this study are available from the corresponding author upon reasonable request.

## Acknowledgments

This work was supported by NIMH (R01MH106507 to V.S.S. and K99MH108720 to K.K.A.C.), and the Brain and Behavior Foundation (NARSAD Young Investigator Award to K.K.A.C.). We appreciate helpful feedback on earlier versions of this manuscript from Loren Frank and Massimo Scanziani.

## Contributions

K.K.A.C. and V.S.S. designed experiments and wrote the manuscript. K.K.A.C. performed all the experiments. T.J.D. and K.K.A.C. set up the LED photometry rig and the dual-site TEMPO rig. K.K.A.C. generated AAV5-I12b-BG-DIO-eYFP. J.D.M. and M.J.S. provided guidance, advice, and feedback on the acquisition and analysis of TEMPO data. K.K.A.C. and V.S.S. analyzed the data.

## Competing interests

The authors declare no competing interests.

## Corresponding author

Correspondence to Vikaas S. Sohal.

## METHODS

Further information and requests for resources and reagents should be directed to and will be fulfilled by Vikaas Sohal.

### Mice

All animal care, procedures, and experiments were conducted in accordance with the NIH guidelines and approved by the Administrative Panels on Laboratory Animal Care at the University of California, San Francisco. Mice were group housed (2-5 siblings) in a temperature-controlled environment (22-24°C), had *ad libitum* access to food and water, and reared in normal lighting conditions (12-h light-dark cycle), until rule shifting experiments began. *Dlx5/6* mice (Wang et al., 2010, Cho et al., 2015) were backcrossed to C57Bl/6 mice for at least 6 generations and then crossed to the *Cre* driver lines: *PV-Cre* (The Jackson Laboratory), *Sst-Cre* (The Jackson Laboratory), and *Ai14* (The Jackson Laboratory). Both male and female adult mice (10-20 weeks at time of experiment) were used in the behavioral experiments. All experiments were done using *Dlx5/6^+/-^* mice and their age-matched *Dlx5/6^+/+^* littermates (crossed to *PV-Cre, Sst-Cre*, and/or *Ai14* lines). All experiments that contained different groups of mice, e.g., *Dlx5/6^+/+^* and *Dlx5/6^+/-^* mice or ChR2-expressing and eYFP-expressing mice, were performed blind to genotype and/or virus injected. This was the case for all experiments except for the experiments shown in Figure 3 (in which all mice were *Dlx5/6^+/+^*, *PV-Cre, Ai14*) and Figures 6a–c (in which all mice were *Dlx5/6^+/-^, PV-Cre* and expressed ChR2).

### Cloning of viral constructs

To produce AAV5-I12b-BG-DIO-eYFP, we introduced MluI and BamHI compatible sticky ends to the DlxI12b-BG sequence with PCR. The pAAV-EF1α-DIO-eYFP (Addgene) was then cut with MluI/BamHI and ligated to the PCR insert to exchange the EF1α promoter for DlxI12b-BG. Virus was packaged by Virovek (Hayward, CA) with serotype AAV5.

To produce AAV1-CAG-DIO-Ace2N-4AA-mNeon, we received pAAV-CAG-DIO-Ace2N-4AA-mNeon from Mark J. Schnitzer (Stanford University). Virus was packaged by Virovek with serotype AAV1.

### Surgery

Male and female mice were anaesthetized using isoflurane (2.5% induction, 1.2-1.5% maintenance, in 95% oxygen) and placed in a stereotaxic frame (David Kopf Instruments). Body temperature was maintained using a heating pad. An incision was made to expose the skull for stereotaxic alignment using bregma and lambda as vertical references. The scalp and periosteum were removed from the dorsal surface of the skull and scored with a scalpel to improve implant adhesion. Viruses were infused at 100-150 nL/min through a 35-gauge, beveled injection needle (World Precision Instruments) using a microsyringe pump (World Precision Instruments, UMP3 UltraMicroPump). After infusion, the needle was kept at the injection site for 5-10 min and then slowly withdrawn. After surgery, mice were allowed to recover until ambulatory on a heated pad, then returned to their homecage.

For behavioral experiments using Cre-dependent optogenetic opsins, mice were injected bilaterally in the mPFC, near the border between the prelimbic and infralimbic cortices (1.7 anterior-posterior (AP), ±0.3 mediolateral (ML), and −2.75 dorsoventral (DV) millimeters relative to bregma) with 1 μL of AAV5-EF1α-DIO-ChR2-eYFP (UNC Virus Core) or 1 μL of AAV5-I12b-BG-DIO-ChR2-eYFP or 1 μL of AAV5-EF1α-DIO-eYFP (UNC VIrus Core) per hemisphere, to selectively target neurons expressing Cre. *Dlx5/6, Sst-Cre* mice were injected bilaterally in the mPFC (1.7 (AP), ±0.3 (ML), and −2.75 (DV)) with 1 μL of AAV5-EF1α-DIO-ChR2-eYFP or 1 μL of AAV5-EF1α-DIO-eYFP per hemisphere. After injection of virus, a 200/240 μm (core/outer) diameter, NA=0.22, dual fiber-optic cannula (Doric Lenses, DFC_200/240-0.22_2.3mm_GS0.7_FLT) was slowly inserted into mPFC until the tip of the fiber reached a DV depth of −2.25. Implants were affixed onto the skull using Metabond Quick Adhesive Cement (Parkell). We waited at least 5 weeks after injection before behavioral experiments to allow for virus expression. For experiments using LFP recordings, standard-tip 0.4 MΩ-impedance tungsten microelectrodes (Microprobes) were used. The coordinates were adjusted to accommodate experiments whereby LFP electrodes were affixed to the fiber implant and protruded 200-300 μm beyond the fiber tip. A common reference screw was implanted into the cerebellum: −5 (AP), 0 (ML) and a ground screw was implanted at −5 (AP), −3 (ML). After affixing the electrodes in place using Metabond (Parkell), connections were made to the headstage of a multi-channel recording system (Pinnacle Technology, Inc.).

For behavioral experiments used in photometry experiments, mice were injected unilaterally at 4 depths (DV: −2.75, −2.5, −2.25, −2.0) at the following AP/ML for mPFC: 1.7 AP, ±0.3 ML with 4 x 0.2 μL of AAV2/1-Syn-FLEX-GCaMP6f-WPRE-SV40 (UPenn Virus Core). After injection of virus, a 400/430 μm (core/outer) diameter, NA=0.48, multimode fiber implant (Doric Lenses, MFC_400/430-0.48_2.8mm_ZF2.5_FLT) was slowly inserted into the mPFC until the tip of the fiber reached a DV depth of −2.25. We waited at least 4 weeks after injection before behavioral experiments to allow for virus expression.

For behavioral experiments used in dual-site TEMPO experiments, mice were injected bilaterally at 3 depths (DV: −2.5, −2.25, −2.0) at the following AP/ML for mPFC: 1.7 AP, ±0.3 (ML) with 3 x 0.2 μL of AAV1-CAG-DIO-Ace2N-4AA-mNeon (Virovek) or with the addition of 0.1 μL per depth of AAV2-Syn-tdTomato (SignaGen Laboratories). After injection of virus, a 400/430 μm (core/outer) diameter, NA=0.48, multimode fiber implant (Doric Lenses, MFC_400/430-0.48_2.8mm_ZF1.25_FLT) was slowly inserted into the mPFC at a 12° angle using the following coordinates: 1.7 (AP), ±0.76 (ML), −2.13 (DV). We waited at least 5 weeks after injection before behavioral experiments to allow for virus expression.

### Rule shift task

This cognitive flexibility task was described in Cho et al., 2015. Briefly, mice are singly-housed and habituated to a reverse light/dark cycle and food intake is restricted until the mouse is 80-85% of the *ad libitum* feeding weight. At the start of each trial, the mouse was placed in its home cage to explore two bowls, each containing one odor and one digging medium, until it dug in one bowl, signifying a choice. As soon as a mouse began to dig in one bowl, the other bowl was removed, so there was no opportunity for “bowl switching.” The bait was a piece of a peanut butter chip (approximately 5-10 mg in weight) and the cues, either olfactory (odor) or somatosensory and visual (texture of the digging medium which hides the bait), were altered and counterbalanced. All cues were presented in small animal food bowls (All Living Things Nibble bowls, PetSmart) that were identical in color and size. Digging media were mixed with the odor (0.01% by volume) and peanut butter chip powder (0.1% by volume). All odors were ground dried spices (McCormick garlic and coriander), and unscented digging media (Mosser Lee White Sand Soil Cover, Natural Integrity Clumping Clay cat litter).

After mice reached their target weight, they underwent one day of habituation. On this day, mice were given ten consecutive trials with the baited food bowl to ascertain that they could reliably dig and that only one bowl contained food reward. All mice were able to dig for the reward. Mice do not undergo any other specific training before being tested on the task. Then, on Days 1 and 2 (and in some cases, on additional days as well), mice performed the task (this was the testing done for experiments). After the task was done for the day, the bowls were filled with different odor-medium combinations and food was evenly distributed among these bowls and given to the mouse so that the mouse would disregard any associations made earlier in the day.

Mice were tested through a series of trials. The determination of which odor and medium to pair and which side (left or right) contained the baited bowl was randomized (subject to the requirement that the same combination of pairing and side did not repeat on more than 3 consecutive trials) using http://random.org. On each trial, while the particular odor-medium combination present in each of the two bowls may have changed, the particular stimulus (e.g., a particular odor or medium) that signaled the presence of food reward remained constant over each portion of the task (initial association and rule shift). If the initial association paired a specific odor with food reward, then the digging medium would be considered the irrelevant dimension. The mouse is considered to have learned the initial association between stimulus and reward if it makes 8 correct choices during 10 consecutive trials. Each portion of the task ended when the mouse met this criterion. Following the initial association, the rule shift portion of the task began, and the particular stimulus associated with reward underwent an extra-dimensional shift. For example, if an odor had been associated with reward during the initial association, then a digging medium was associated with reward during the rule shift portion of the task. The mouse is considered to have learned this extra-dimensional rule shift if it makes 8 correct choices during 10 consecutive trials. When a mouse makes a correct choice on a trial, it is allowed to consume the food reward before the next trial. Following correct trials, the mouse is transferred from the home cage to a holding cage for about 10 seconds while the new bowls were set up (intertrial interval). After making an error on a trial, a mouse was transferred to the holding cage for about 2 minutes (intertrial interval). All animals performed the initial association in a similar number of trials (average: 10-15 trials). We were blind to genotype and/or virus injected.

### Rule reversal task

This cognitive flexibility task was described in Cho et al., 2015. Similar to the mechanics of the rule-shift task described above, following the initial association, the rule reversal portion of the task began, and the particular stimulus associated with reward underwent an intra-dimensional shift. For example, if an odor had been associated with reward during the initial association, then the previously uncued odor was associated with reward during the rule reversal portion of the task. The mouse is considered to have learned this intra-dimensional rule reversal if it makes 8 correct choices during 10 consecutive trials.

Mice that were involved in both the rule shift and rule reversal tasks were randomly assigned the order of tasks over the course of two days.

### *In vivo* optogenetic stimulation

In-phase ChR2 stimulation: A 473 nm blue laser (OEM Laser Systems, Inc.) was coupled to the dual fiber-optic cannula (Doric Lenses) through a 200 μm diameter dual fiber-optic patchcord with guiding socket (Doric Lenses, Inc.) and 1×2 intensity division fiber-optic rotary joint (Doric Lenses, Inc.), and adjusted such that the final light power was ~0.5 mW total, summed across both fibers and averaged over light pulses and the intervening periods. A function generator (Agilent 33500B Series Waveform Generator) connected to the laser generated a 40 Hz train of 5 ms pulses.

Out-of-phase ChR2 stimulation: The stereotaxically implanted dual fiber-optic cannula was coupled to two separate 473 nm blue lasers via a dual fiber-optic patch cord with fully-separated optical paths that were each connected to separate fiber-optic rotary joints. Again, light power was adjusted such that the final output was ~0.5 mW across both fibers. Different function generators connected to each laser, in order to generate out-of-phase stimulation. For the experiments shown in Figures 4d–e and Figures 6d–f, these two function generators were not connected in any way, except that we verified (by eye) that the light pulses were delivered at non-overlapping times, producing phase differences between 72 and 288 degrees. For the experiments shown in Figures 4a–c and Figures 4f–h, one function generator was triggered at the time when the other function generator switched off, so the phase difference was exactly 72 degrees. Stimulation was generated using either a 20 Hz train of 10ms pulses or a 40 Hz train of 5ms pulses.

For all experiments in which we delivered optogenetic stimulation to behaving mice, light stimulation began once mice reached the 80% criterion during the initial association portion of the task. Mice then performed three additional initial association trials with the light stimulation before the rule shift portion of the task began. The light stimulation did not alter the performance or behavior of the mice during these three extra trials of the initial association. Experiments were performed blind to genotype and/or virus injected.

### Drug administration

Clonazepam at indicated concentrations (0.0625 mg/kg, Sigma) was diluted in the vehicle solution (PBS with 0.5% methylcellulose) then injected (I.P.) in a volume of 0.01 ml/kg 30 min prior to behavioral testing.

### Fiber photometry design and recording

The photometry apparatus and analysis was based on Lerner et al., 2015, with some modifications described below.

A fiber-optic stub (400 μm core, NA=0.48, low-autofluorescence fiber; Doric Lenses, Quebec, Canada, MFC_400/430-0.48_2.3mm_ZF2.5_FLT) was stereotaxically implanted in mPFC. A single fiber was used to both deliver excitation light and collect emitted fluorescence from the recording site. A matching fiberoptic patch cord (Doric Lenses, MFP_400/430/1100-0.48_2m_FC-ZF2.5) provided a light path between the animal and a miniature, permanently-aligned optical bench, or ‘mini cube’ (Doric Lenses, FMC2_AF405-GCaMP_FC). Two excitation LEDs (470 nm ‘blue’ and 405 nm ‘violet’, Thorlabs M470F1 and M405FP1) were connected to the ‘mini cube’ by a patch cord (200 μm core, NA = 0.39, Doric Lenses) and controlled by an LED driver (Thorlabs DC4104), and connected to an RX-8 real-time processor (Tucker Davis Technologies). Excitation light is delivered at 470 nm to stimulate GCaMP6f fluorescence in a Ca^2+^-dependent manner and at 405 nm, an excitation isosbestic wavelength for GCaMP6f, to perform ratiometric measurements of GCaMP6f activity, correcting for bleaching and artifactual signal fluctuations. Blue excitation was sinusoidally-modulated at 210 Hz and violet excitation was modulated at 330 Hz. The GCaMP6f emission signal was collected through a patchcord (Doric Lenses, MFP_600/630/LWMJ-0.48_0.5m_FC-FC) and focused onto a femtowatt photoreceiver (Newport, Model 2151) with a lensed, permanently-aligned FC coupler (Doric Lenses). Each of the two modulated signals generated by the two LEDs was independently recovered using standard synchronous demodulation techniques implemented on the RX-8 real-time processor. The commercial software Synapse (Tucker-Davis Technologies) running on a PC was used to control the signal processor, write data streams to disk, and to record synchronized video from a generic infrared USB webcam (Ailipu Technology, Shenzhen, China, ELP-USB100W05MT-DL36). Files were then exported for analysis to MATLAB (Mathworks).

For every experiment, the far end of the patch cord and the 2.5 mm diameter zirconia optical implant ferrule were cleaned with isopropanol before each recording, then securely attached via a zirconia sleeve.

### LFP recording

Data was recorded at 2 kHz and band-pass filtered at 0.5-200 Hz at the pre-amplifier from mice using a time-locked video multi-channel recording system (Pinnacle Technology, Inc.) using Sirenia Acquisition software (Pinnacle Technology, Inc.). Channels shared a common reference (cerebellum). Mice were recorded in their home cage, including a baseline and stimulation of a 40 Hz train of 5ms pulses with a 473 nm blue laser (OEM Laser Systems, Inc.).

### Dual-site voltage-sensor photometry (TEMPO)

High-bandwidth bandwidth time-varying bulk fluorescence signals were measured at each recording site using the TEMPO technique described in Marshall et al., 2016, with some modifications as described below.

### Optical apparatus

A fiber-optic stub (400 μm core, NA=0.48, low-autofluorescence fiber; Doric Lenses, MFC_400/430-0.48_2.8mm_ZF1.25_FLT) was stereotaxically implanted in each targeted brain region. A matching fiberoptic patch cord (Doric Lenses, MFP_400/430/1100-0.48_2m_FC-ZF1.25) provided a light path between the animal and a miniature, permanently-aligned optical bench, or ‘mini-cube’ (Doric Lenses, FMC5_E1(460-490)_F1 (500-540)_E2(555-570)_F2(580-680)_S). A single fiber was used to both deliver excitation light to and collect emitted fluorescence from each recording site. The far end of the patch cord and each 1.25mm diameter zirconia optical implant ferrule were cleaned with isopropanol before each recording, then securely attached via a zirconia sleeve.

The mini-cube optics allow for the simultaneous monitoring of two spectrally-separated fluorophores, with dichroic mirrors and cleanup filters chosen to match the excitation and emission spectra of the voltage sensor and reference fluorophores in use (‘mNeon’ voltage sensor channel: Ex. 460-490 nm, Em. 500-540 nm; ‘Red’ control fluorophore: Ex. 555-570 nm, Em. 580-680 nm). The mini-cube optics are sealed and permanently aligned and all 5 ports (sample to animal, 2 excitation lines, 2 emission lines) are provided with matched coupling optics and FC connectors to allow for a modular system design.

Excitation light for each of the two color channels was provided by a fiber-coupled LED (Center wavelengths 490 nm and 565 nm, Thorlabs M490F3 and M565F3) connected to the mini-cube by a patch cord (200 μm, NA=0.39; Thorlabs M75L01). Using a smaller diameter for this patch cord than for the patch cord from the cube to the animal is critical to reduce the excitation spot size on the output fiber face and thus avoid cladding autofluorescence. LEDs were controlled by a 4-channel, 10kHz-bandwidth current source (Thorlabs DC4104). LED current was adjusted to give a final light power at the animal (averaged during modulation, see below) of approximately 200 μW for the mNeon channel (460-490 nm excitation), and 100 μW for the Red channel (555-570 nm excitation).

Each of the two emission ports on the mini-cube was connected to an adjustable-gain photoreceiver (Femto, Berlin, Germany, OE-200-Si-FC; Bandwidth set to 7kHz, AC-coupled, ‘Low’gain of ~5×10^7 V/W) using a large-core high-NA fiber to maximize throughput (600 μm core, NA=0.48 (Doric lenses, MFP_600/630/LWMJ-0.48_0.5m_FC-FC).

Note that, for dual-site recordings, two completely independent optical setups were employed, with separate implants, patch cords, mini-cubes, LEDs, photoreceivers, and lock-in amplifiers.

### Modulation and lock-in detection

At each recording site, each of the two LEDs was sinusoidally modulated at a distinct carrier frequency to reduce crosstalk due to overlap in fluorophore spectra. The corresponding photoreceiver outputs were then demodulated using lock-in amplification techniques. A single instrument (Stanford Research Systems, SR860) was used to generate the modulation waveform for each LED and to demodulate the photoreceiver output at the carrier frequency. To further reduce cross-talk between recording sites, distinct carrier frequencies were used across at each site (mNeon site 1: 2 kHz; mNeon site 2: 2.5 kHz; Red site 1: 3.5 kHz; Red site 2: 4 kHz). Low-pass filters on the lock-in amplifiers were selected to reject noise above the frequencies under study (cascade of 4 Gaussian FIR filters with 84 Hz equivalent noise bandwidth; final attenuation of signals are approximately −1dB (89% of original magnitude) at 20 Hz, −3dB (71% of original magnitude) at 40 Hz, and −6dB (50% of original magnitude) at 60 Hz).

### TEMPO recording

Analog signals were digitized by a multichannel real-time signal processor (Tucker-Davis Technologies, Alachua, Florida; RX-8). The commercial software Synapse (Tucker-Davis Technologies) running on a PC was used to control the signal processor, write data streams to disk, and to record synchronized video from a generic infrared USB webcam (Ailipu Technology, Shenzhen, China, ELP-USB100W05MT-DL36). Lock-in amplifier outputs were digitized at 3 kHz.

### Histology and imaging

All mice used for behavioral and imaging experiments were anesthetized with Euthasol and transcardially perfused with 30 mL of ice-cold 0.01 M PBS followed by 30 ml of ice-cold 4% paraformaldehyde (PFA) in PBS. Brains were extracted and stored in 4% PFA for 24 h at 4 °C before being stored in PBS. 50 μm and 70 μm slices were obtained on a Leica VT100S and mounted on slides. All imaging was performed on an Olympus MVX10, Nikon Eclipse 90i, Zeiss LSM510, and Zeiss Axioskop2. We verified that all mice had virus-driven expression and optical fibers located in the mPFC.

### Fiber photometry analysis

For calculating peak GCaMP6f signals during activity time-locked to task-related events: the beginning of each trial, the time of each decision (indicated by the mouse beginning to dig in one bowl), and the beginning and end of each intertrial interval, a least-squares linear fit was applied to the 405 nm control signal to align it to the 470 nm signal. The *dF/F* time series was then calculated for each behavioral session as: ((470 nm signal - fitted 405 nm signal) / fitted 405 nm signal). For points of interest (e.g., time of decision), peak *dF/F* was calculated as the most extreme *dF/F* value at time 0-4 seconds (positive or negative). Experiments were performed, scored, and analyzed blind to genotype.

### LFP analysis

To analyze changes in power in our LFP data, we computed spectrograms from completely nonoverlapping time windows and compared unnormalized power (measured in log units) during 40 Hz optical stimulation to the power during the baseline period.

### TEMPO analysis

Analysis of TEMPO data was facilitated using the signal processing toolbox and MATLAB (Mathworks), using the following functions: fir1, filtfilt, and regress. All four signals during the entire time series of the experiment (left mNeon, left tdTomato, right mNeon, right tdTomato) were first filtered around a frequency of interest. To quantify zero-phase lag interhemispheric synchronization between left and right mNeon signals, we performed a linear regression analysis to predict the right mNeon signal using the following inputs: left mNeon signal, left tdTomato signal, and right tdTomato signal. The goodness of fit is compared to how well the regression works if the left mNeon signal is shuffled, i.e., if we use a randomly chosen segment of the original left mNeon signal, instead of the segment recorded at the same time as the right mNeon signal. *R^2^* values are calculated as a function of time and compared to the 99^th^ percentile of the distribution of *R^2^* values obtained from 100 fits to randomly-shuffled data. The fraction of timepoints at which the *R^2^* obtained from actual data exceeds the 99^th^ percentile of the *R^2^* values obtained from shuffled data was used to measure zero-phase lag synchronization between the left and right mNeon signals.

We performed this analysis at the time of the decision (e.g., immediately following the beginning of digging in one bowl, until the end of digging). To measure 90 degree out-of-phase synchronization, the filtered left mNeon signal was simply advanced 90 degrees relative to the right mNeon signal, before performing the analysis described above. Experiments were performed, scored, and analyzed blind to genotype. Mice that did not make both correct and incorrect decisions in the first 5 trials of a task were excluded from analyses involving comparisons of activity on correct vs. incorrect decisions.

### Data analyses and statistics

Statistical analyses were performed using Prism 7 (GraphPad) and detailed in the corresponding figure legends. Quantitative data are expressed as the mean and error bars or shaded areas represent the standard error of the mean (s.e.m.). Group comparisons were made using one-way repeated measures or two-way ANOVA followed by Tukey’s or Bonferroni post-hoc tests to control for multiple comparisons, respectively. Paired and unpaired two-tailed Student’s *t*-tests were used to make single-variable comparisons or with Welch’s correction for unequal variance. Similarity of variance between groups was confirmed by the *F* test. Measurements were taken from distinct samples. *P* = *, < 0.05; **, < 0.01; ***, < 0.001; ****, < 0.0001. Comparisons with no asterisk had *P* > 0.05 and were considered not significant.

