## Supplemental Figures for "Interhemispheric gamma synchrony between parvalbumin interneurons supports behavioral adaptation"

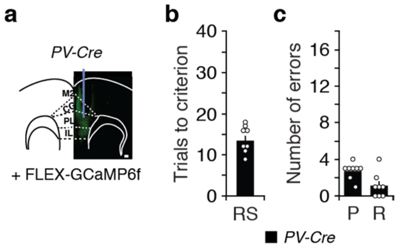


**Supplementary Fig. 1: Learning during rule shifts in mice used for photometry experiments.**

**a**, *PV-Cre* mice had a unilateral FLEX-GCaMP6f injection and fiber-optic implant in mPFC for photometry (scale bar, 100 μm).
**b**, Rule-shift (RS) performance of *PV-Cre* mice ($n$ = 8) used for photometry experiments.
**c**, Numbers of perseverative (P) or random (R) errors during the rule shift.
Data are shown as means (**b**, **c**); error bars (**b**, **c**) denote s.e.m.

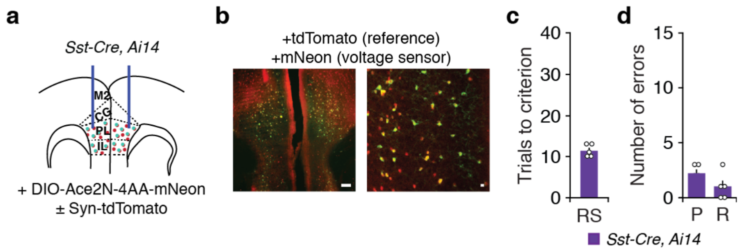


**Supplementary Fig. 2: Learning during rule shifts in mice used for TEMPO imaging from Sst interneurons.**

**a**, *Sst-Cre, Ai14* mice (*n* = 5) had bilateral AAV-DIO-Ace2N-4AA-mNeon ± AAV-Syn-tdTomato injections and fiber-optic implants in mPFC.
**b**, Examples of tdTomato (red) and mNeon (green) fluorescence in a coronal section of mPFC (left), alongside a high power image (right). Scale bars: 100 μm and 25 μm, respectively.
**c**, Rule-shift (RS) performance of *Sst-Cre, Ai14* mice (*n* = 5) used for dual-site TEMPO imaging.
**d**, Number of perseverative (P) and random (R) errors during the rule shift.
Data are shown as means (**c**, **d**); error bars (**c**, **d**) denote s.e.m.

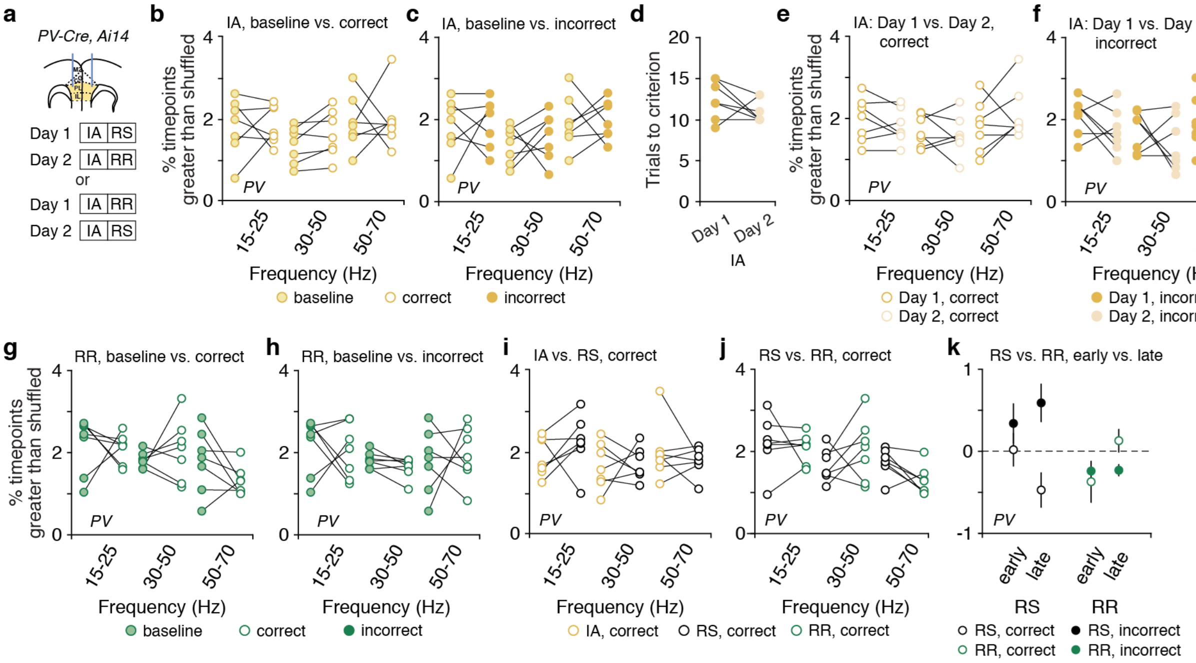


**Supplementary Fig. 3: Interhemispheric synchrony between prefrontal PV interneurons for various frequency bands and types of trials.**

**a**, *PV-Cre, Ai14* mice ($n$ = 7) had bilateral AAV-DIO-Ace2N-4AA-mNeon ± AAV-Syn-tdTomato injections and fiber-optic implants in mPFC. Experimental design: Day 1: Initial association (IA) followed by rule shift (RS) or rule reversal (RR); Day 2: IA followed by the rule change (RS or RR) that was not performed on Day 1.

**b**, During learning of the IA that preceded the RS, synchrony was not different after correct decisions vs. during the baseline period (two-way ANOVA; main effect of condition: $F_{1,18}$ = 0.51, $P$ = 0.48; frequency X condition interaction: $F_{2,18}$ = 0.20, $P$ = 0.82).

**c**, During learning of this IA, synchrony was not different after incorrect decisions vs. during the baseline period (two-way ANOVA; main effect of condition: $F_{1,18}$ = 0.39, $P$ = 0.54; frequency X condition interaction: $F_{2,18}$ = 0.07, $P$ = 0.94).

**d**, IA performance was not different across days (two-tailed, paired $t$-test; $t_{(6)}$ = 1.29, $P$ = 0.25).

**e**, There was no difference in synchrony after correct trials during learning of the IA on Day 1 vs. 2 (two-way ANOVA; main effect of day: $F_{1,18}$ = 0.02, $P$ = 0.89; frequency X condition interaction: $F_{2,18}$ = 1.48, $P$ = 0.26).

**f**, There was no difference in synchrony after incorrect trials during learning of the IA on Day 1 vs. 2 (two-way ANOVA; main effect of day: $F_{1,18}$ = 3.05, $P$ = 0.10; frequency X condition interaction: $F_{2,18}$ = 0.03, $P$ = 0.97).

**g**, During the RR, synchrony was not different after correct decisions vs. during the baseline period (two-way ANOVA; main effect of condition: $F_{1,18}$ = 0.28, $P$ = 0.60; frequency X condition interaction: $F_{2,18}$ = 1.24, $P$ = 0.31).

**h**, During the RR, synchrony was not different after incorrect decisions vs. the baseline period (two-way ANOVA; main effect of condition: $F_{1,18}$ = 0.07, $P$ = 0.79; frequency X condition interaction: $F_{2,18}$ = 0.28, $P$ = 0.76).

**i**, Synchrony after correct decisions did not differ between the IA vs. RS (two-way ANOVA; main effect of condition: $F_{1,18}$ = 0.13, $P$ = 0.73; frequency X condition interaction: $F_{2,18}$ = 1, $P$ = 0.39).

**j**, Synchrony after correct decisions did not differ between the RR vs. RS (two-way ANOVA; main effect of condition: $F_{1,18}$ = 0.16, $P$ = 0.70; frequency X condition interaction: $F_{2,18}$ = 2.55, $P$ = 0.11).

**k**, The plot shows the average gamma synchrony on correct vs. incorrect trials, during the first 2 ('early') or next 3 ('late') trials of a rule shift (RS) or rule reversal (RR). In order to average together values from different mice, each synchrony value was computed relative to the average gamma synchrony measured during the first 5 RS and RR trials from the same mouse. We

performed ANOVA on the gamma synchrony from each of the first 5 trials during a rule shift (RS) or rule reversal (RR), including the following factors and interaction terms: mouse ($F_{6,59}$ = 1.72, *P* = 0.13), type of rule change (RS vs. RR) ($F_{1,59}$ = 3.74, *P* = 0.06), correct vs. incorrect trial outcome ($F_{1,59}$ = 2.64, *P* = 0.11), an interaction of correct-incorrect X RS-RR ($F_{1,59}$ = 11.12, ***P* = 0.0015), and an interaction of correct-incorrect X RS-RR X early vs. late trials (i.e. first 2 vs. next 3 trials) ($F_{1,59}$ = 4.28, **P* = 0.043).
Two-way ANOVA followed by Bonferroni post hoc comparisons were used in panels **b**–**c** and **e**–**j**. Comparisons were not significant unless otherwise noted.

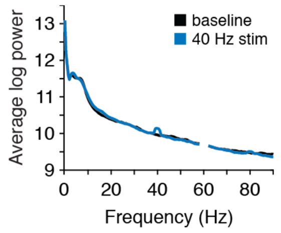


**Supplementary Fig. 4: Prefrontal LFP power spectrum with or without 40 Hz optogenetic stimulation of PV interneurons.**

Average log power for prefrontal LFP recordings ipsilateral to stimulation from *PV-Cre* mice (*n* = 3) during 40 Hz stimulation (blue trace) and baseline (black trace). Data are shown as means.

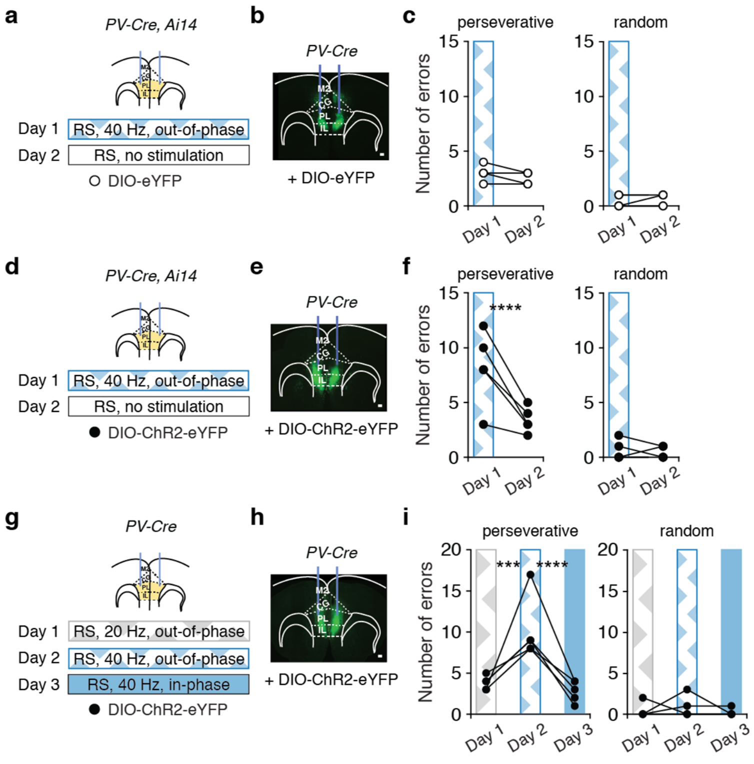


**Supplementary Fig. 5. Types of errors during rule shifts (RS) in the presence of various types of optogenetic stimulation.**
**a, d**, *PV-Cre, Ai14* mice had bilateral AAV-DIO-eYFP (**a**; $n$ = 5) or AAV-DIO-ChR2 (**d**; $n$ = 5) injections and fiber-optic implants in mPFC. Experimental design: Day 1: out-of-phase 40 Hz stimulation during the rule shift (RS); Day 2: no stimulation.
**b, e**, Representative images showing mPFC expression of eYFP (**b**) or ChR2-eYFP (**e**) (scale bar, 100 µm).
**c, f,** Optogenetic stimulation increases perseverative errors in ChR2-expressing mice compared to eYFP-expressing controls (two-way ANOVA; main effect of day: $F_{1,8}$ = 35.9, ***$P$ = 0.0003; main effect of virus: $F_{1,8}$ = 46.9, ***$P$ = 0.0001; day X virus interaction: $F_{1,8}$ = 30.5, ***$P$ = 0.0006). There is no change in random errors (two-way ANOVA; main effect of day: $F_{1,8}$ = 0, $P$ > 0.99; main effect of virus: $F_{1,8}$ = 0, $P$ > 0.99; day X virus interaction: $F_{1,8}$ = 0.89, $P$ = 0.37).
**c**, Light delivery does not affect the number of perseverative or random errors in eYFP-expressing controls (perseverative: post hoc $t_{(8)}$ = 0.33, $P$ > 0.99; random: post hoc $t_{(8)}$ = 0.67, $P$ > 0.99).
**f**, Optogenetic stimulation of PV interneurons on Day 1 increased the number of perseverative errors compared to no stimulation on Day 2 (post hoc $t_{(8)}$ = 8.14, ****$P$ < 0.0001), but does not affect random errors (post hoc $t_{(8)}$ = 0.67, $P$ > 0.99).
**g**, *PV-Cre* mice ($n$ = 5) had bilateral AAV-DIO-ChR2-eYFP injections and fiber-optic implants in mPFC. Experimental design: Day 1: out-of-phase 20 Hz stimulation; Day 2: out-of-phase 40 Hz stimulation; Day 3: in-phase 40 Hz stimulation.
**h**, Representative image showing mPFC ChR2 expression (scale bar, 100 µm).
**i**, Out-of-phase 40 Hz stimulation (Day 2) increases perseverative errors but does not affect random errors, relative to out-of-phase 20 Hz stimulation (Day 1) or in-phase 40 Hz stimulation (Day 3) (two-way ANOVA; main effect of day: $F_{2,16}$ = 13.7, ***$P$ = 0.0003; Day 1 vs. Day 2 perseverative: ***$P$ = 0.0002, Day 2 vs. Day 3 perseverative: ****$P$ < 0.0001, Day 1 vs. Day 3 perseverative: $P$ > 0.99; Day 1 vs. Day 2 random: $P$ > 0.99, Day 2 vs. Day 3 random: $P$ > 0.99, Day 1 vs. Day 3 random: $P$ > 0.99).
Two-way ANOVA followed by Bonferroni post hoc comparisons were used.

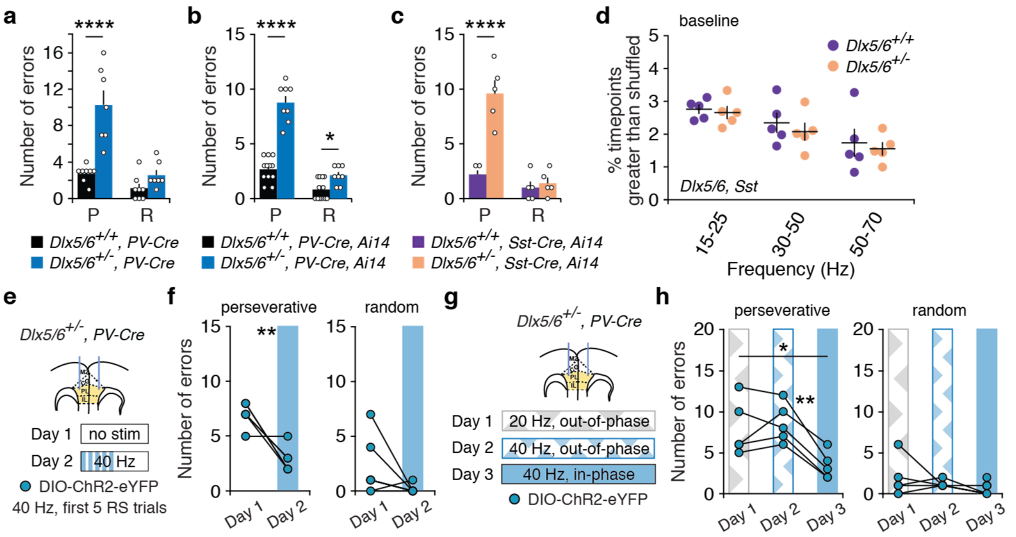


**Supplementary Fig. 6: Types of errors during rule shifts in *Dlx5/6*$^{+/-}$ and wild-type mice.**

**a**, Rule-shift (RS) performance for photometry experiments in *Dlx5/6*$^{+/-}$*, PV-Cre* and *Dlx5/6*$^{+/+}$*, PV-Cre* mice. Compared to wild-type littermates, mutant mice make more perseverative errors (two-way ANOVA; main effect of genotype: $F_{1,13}$ = 39.3, ****$P$ < 0.0001; main effect of error type: $F_{1,13}$ = 25.3, ***$P$ < 0.001; error type X genotype interaction: $F_{1,13}$ = 10.8, **$P$ = 0.006; post hoc $t_{(26)}$ = 6.43, ****$P$ < 0.0001), but similar numbers of random errors (post hoc $t_{(26)}$ = 1.23, $P$ = 0.46).

**b**, Rule-shift performance for mice used in dual-site TEMPO experiments. Compared to wild-type mice, mutant mice make more perseverative errors (two-way ANOVA; main effect of genotype: $F_{1,18}$ = 84.1, ****$P$ < 0.0001; main effect of error type: $F_{1,13}$ = 148.8, ****$P$ < 0.0001; type of error X genotype interaction: $F_{1,18}$ = 47.8, ****$P$ < 0.0001; post hoc $t_{(36)}$ = 11.5, ****$P$ < 0.0001), and random errors (post hoc $t_{(36)}$ = 2.43, *$P$ = 0.04).

**c**, Compared to wild-type mice, *Dlx5/6*$^{+/-}$*, Sst-Cre, Ai14* mice make more perseverative errors (two-way ANOVA; main effect of genotype: $F_{1,8}$ = 42.3, ***$P$ = 0.0002; main effect of error type: $F_{1,13}$ = 30.7, ***$P$ < 0.001; error type X genotype interaction: $F_{1,8}$ = 17.0, **$P$ = 0.003; post hoc $t_{(16)}$ = 7.12, ****$P$ < 0.0001), but numbers of random errors are comparable (post hoc $t_{(16)}$ = 0.38, $P$ > 0.99).

**d**, Synchrony was not different between *Dlx5/6*$^{++-}$*, Sst-Cre, Ai14* and *Dlx5/6*$^{+/-}$*, Sst-Cre, Ai14* mice during the baseline period (two-way ANOVA; main effect of genotype: $F_{1,12}$ = 0.55, $P$ = 0.47; frequency X genotype interaction: $F_{2,12}$ = 0.04, $P$ = 0.96).

**e**, *Dlx5/6*$^{+/-}$, *PV-Cre* mice ($n$ = 6) had bilateral AAV-DIO-ChR2-eYFP injections and fiber-optic implants in mPFC. Experimental design: Day 1: no stimulation; Day 2: in-phase 40 Hz stimulation during the first 5 RS trials.

**f**, In-phase 40 Hz stimulation on Day 2 reduces perseverative errors relative to no stimulation on Day 1 (two-way ANOVA; main effect of day: $F_{1,10}$ = 49.9, ****$P$ < 0.0001; main effect of error type: $F_{1,10}$ = 18.3, **$P$ = 0.0016; post hoc $t_{(10)}$ = 3.98, **$P$ = 0.005); there was no change in random errors (post hoc $t_{(10)}$ = 2.07, $P$ = 0.13).

**g**, *Dlx5/6*$^{+/-}$, *PV-Cre* mice ($n$ = 5) had bilateral AAV-DIO-ChR2-eYFP injections and fiber-optic implants in mPFC. Experimental design: Day 1: out-of-phase 20 Hz stimulation; Day 2: out-of-phase 40 Hz stimulation; Day 3: in-phase 40 Hz stimulation.

**h**, Perseverative errors are reduced by in phase 40 Hz stimulation on Day 3, compared to either out-of-phase 20 Hz stimulation on Day 1 or out-of-phase 40 Hz stimulation on Day 2 (two-way ANOVA; main effect of day: $F_{2,16}$ = 12.9, ****$P$ = 0.0005; main effect of error type: $F_{1,8}$ = 29.1, ***$P$ = 0.0006; Day 1 vs. Day 2 perseverative: $P$ > 0.99, Day 2 vs. Day 3 perseverative: ***$P$ = 0.0003, Day 1 vs. Day 3 perseverative: ***$P$ = 0.001). There are no changes in random errors across days (post hoc Day 1 vs. Day 2: $P$ > 0.99, Day 2 vs. Day 3: $P$ > 0.99, Day 1 vs. Day 3: $P$ = 0.28). Two-way ANOVA followed by Bonferroni post hoc comparisons were used. Comparisons were not significant unless otherwise noted.

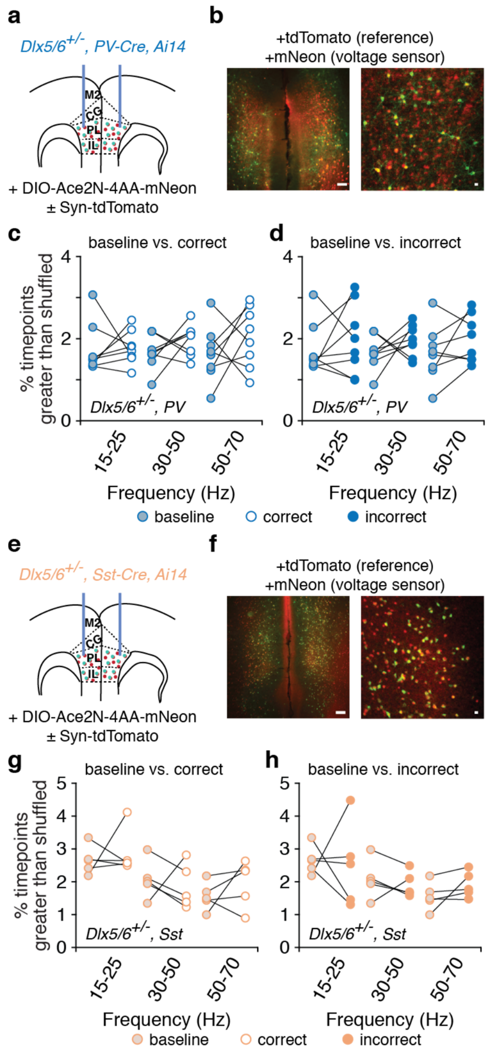


**Supplementary Fig. 7. Interhemispheric synchrony between prefrontal PV or Sst interneurons during rule shifts in *Dlx5/6*$^{+/-}$ mice.**

**a**, *Dlx5/6*$^{+/-}$*, PV-Cre, Ai14* mice (*n* = 8) had bilateral AAV-DIO-Ace2N-4AA-mNeon ± AAV-Syn-tdTomato injections and fiber-optic implants in mPFC.

**b**, Examples of tdTomato (red) and mNeon (green) fluorescence within PV interneurons in a coronal section of mPFC (left), alongside a high power image (right). Scale bars: 100 μm and 25 μm, respectively.

**c**, In mutants, PV interneuron synchrony was not different after correct decisions vs. during the baseline period (two-way ANOVA; main effect of condition: $F_{1,21}$ = 1.89, $P$ = 0.18; frequency X condition interaction: $F_{2,21}$ = 0.35, $P$ = 0.71).

**d**, PV interneuron synchrony was not different after incorrect decisions vs. during the baseline period (two-way ANOVA; main effect of condition: $F_{1,21}$ = 3.31, $P$ = 0.083; frequency X condition interaction: $F_{2,21}$ = 0.04, $P$ = 0.96).

**e**, *Dlx5/6*$^{+/-}$*, Sst-Cre, Ai14* mice (*n* = 5) had bilateral AAV-DIO-Ace2N-4AA-mNeon ± AAV-Syn-tdTomato injections and fiber-optic implants in mPFC.

**f**, Examples of tdTomato (red) and mNeon (green) fluorescence in Sst interneurons within a coronal section of mPFC (left) from a *Dlx5/6*$^{+/-}$*, Sst-Cre, Ai14* mouse, alongside a high power image (right). Scale bars: 100 μm and 25 μm, respectively.

**g**, In mutants, Sst interneuron synchrony was not different after correct decisions vs. during the baseline period (two-way ANOVA; main effect of frequency: $F_{1,21}$ = 6.07, $P$ = 0.015; frequency X condition interaction: $F_{2,12}$ = 0.10, $P$ = 0.90).

**h**, Sst interneuron synchrony was not different after incorrect decisions vs. during the baseline period (two-way ANOVA; main effect of frequency: $F_{1,21}$ = 7.34, $P$ = 0.0083; frequency X condition interaction: $F_{2,12}$ = 0.26, $P$ = 0.78).
Two-way ANOVA followed by Bonferroni post hoc comparisons were used. Comparisons were not significant unless otherwise noted.

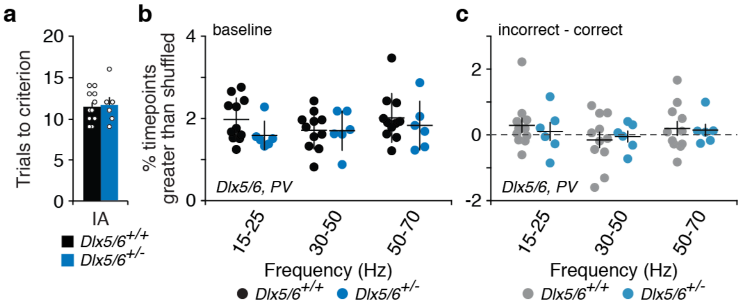


**Supplementary Fig. 8. Interhemispheric PV interneuron synchrony in *Dlx5/6*$^{+/-}$ mice vs. wild-type littermates during baseline periods or learning of initial associations.**

**a**, Learning of an initial association (IA) was similar in mutants ($n$ = 6) and their wild-type littermates (two-tailed, unpaired $t$-test; $n$ = 11; $t_{(15)}$ = 0.202, $P$ = 0.842).

**b**, There was no difference in interhemispheric PV interneuron synchronization between mutant and wild-type littermates at baseline (two-way ANOVA; main effect of genotype: $F_{1,15}$ = 1.56, P = 0.23; genotype X frequency interaction $F_{2,30}$ = 0.55, $P$ = 0.58).

**c**, During learning of an initial association, changes in PV interneuron synchrony following errors (relative to synchrony after correct decisions) is similar in mutants and their wild-type littermates (two-way ANOVA; main effect of genotype: $F_{1,15}$ = 0.07, $P$ = 0.80; genotype X frequency interaction: $F_{2,30}$ = 0.16, $P$ = 0.86).

Data are shown as means (**a**–**c**); error bars (**a**–**c**) denote s.e.m. Two-way ANOVA followed by Bonferroni post hoc comparisons were used, unless otherwise noted. Comparisons were not significant unless otherwise noted.

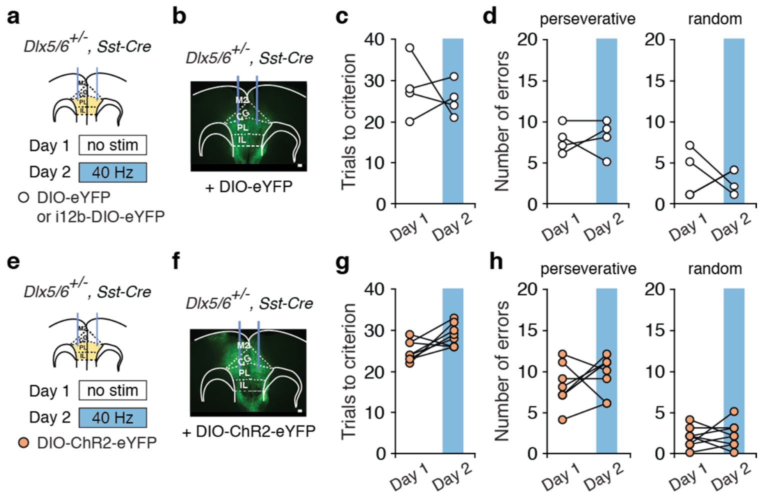


**Supplementary Fig. 9: In-phase, gamma-frequency stimulation of Sst interneurons does not rescue rule shifts in *Dlx5/6*$^{+/-}$, *Sst-Cre* mice.**

**a**,**e,** *Dlx5/6*$^{+/-}$*, Sst-Cre* mice had bilateral control virus (AAV-DIO-eYFP or AAV-i12b-DIO-eYFP; *n* = 4) or AAV-DIO-ChR2-eYFP (**e,** *n* = 8) injections and fiber-optic implants in mPFC. Experimental design: Day 1: no stimulation; Day 2: 40 Hz stimulation.

**b,f**, eYFP (**b**) and ChR2-eYFP (**f**) expression in the mPFC of *Dlx5/6*$^{+/-}$*, Sst-Cre* mice (scale bar, 100 μm).

**c,g,** Light delivery did not affect performance in Sst-eYFP-expressing (**c**) or Sst-ChR2-expressing (**g**) mutant mice (two-way ANOVA; main effect of day: $F_{1,10}$ = 0.08, $P$ = 0.79; main effect of virus: $F_{1,10}$ = 0.002, $P$ = 0.96; day X virus interaction: $F_{1,10}$ = 2.69, $P$ = 0.13).

**d**, Sst-eYFP expressing mutants showed no change in perseverative or random errors from Day 1 to Day 2 (two-way ANOVA; day X type of error interaction: $F_{1,6}$ = 0.25, $P$ = 0.63).

**h**, Sst-ChR2 expressing mutants showed no change in perseverative or random errors from Day 1 to Day 2 (two-way ANOVA; day X type of error interaction: $F_{1,14}$ = 0.44, $P$ = 0.52).

Two-way ANOVA followed by Bonferroni post hoc comparisons were used. Comparisons were not significant unless otherwise noted.
